# Utility of a high-resolution mouse single nucleotide polymorphism microarray assessed for rodent comparative genomics

**DOI:** 10.1101/2020.09.29.318071

**Authors:** Rachel D. Kelly, Maja Milojevic, Freda Qi, Kathleen A. Hill

**Affiliations:** Department of Biology, *The* University *of* Western Ontario, London, Ontario, Canada

## Abstract

In the study of genetic diversity in non-model species there is a notable lack of the low-cost, high resolution tools that are readily available for model organisms. Genotyping microarray technology for model organisms is well-developed, affordable, and potentially adaptable for cross-species hybridization. The Mouse Diversity Genotyping Array (MDGA), a single nucleotide polymorphism (SNP) genotyping tool designed for *Mus musculus*, was tested as a tool to survey genomic diversity of wild species for inter-order, inter-genus, and intra-genus comparisons. Application of the MDGA cross-species provides genetic distance information that reflects known taxonomic relationships reported previously between non-model species, but there is an underestimation of genetic diversity for non-Mus samples, indicated by a plateau in loci genotyped beginning 10-15 millions of years divergence from the house mouse. The number and types of samples included in datasets genotyped together must be considered in cross-species hybridization studies. The number of loci with heterozygous genotypes mapped to published genome sequences indicates potential for cross-species MDGA utility. A case study of seven deer mice yielded 159,797 loci (32% of loci queried by the MDGA) that were genotyped in these rodents. For one species, *Peromyscus maniculatus*, 6,075 potential polymorphic loci were identified. Cross-species utility of the MDGA provides needed genetic information for non-model species that are lacking genomic resources. Genotyping arrays are widely available, developed tools that are capable of capturing large amounts of genetic information in a single application, and represent a unique opportunity to identify genomic variation in closely related species that currently have a paucity of genomic information available. A candidate list of MDGA loci that can be utilized in cross-species hybridization studies was identified and may prove to be informative for rodent species that are known as environmental sentinels. Future studies may evaluate the utility of candidate SNP loci in populations of non-model rodents.

**Author Summary:** There is a need for a tool that can assay DNA sequence differences in species for which there is little or no DNA information available. One method of analyzing differences in DNA sequences in species with well-understood genomes is through a genotyping microarray, which has demonstrated utility cross-species. The Mouse Diversity Genotyping Array (MDGA) is a tool designed to examine known differences across the genome of the house mouse, *Mus musculus*. Given that related organisms share genetic similarity, the MDGA was tested for utility in identifying genome variation in other wild mice and rodents. Variation identified from distantly related species that were not of the same genus as the house mouse was an underestimate of the true amount of variation present in the genomes of wild species. Utility of the MDGA for wild species is best suited to mice from the same genus as the house mouse, and candidate variation identified can be tested in rodent populations in future studies. Identifying changes in genetic variation within populations of wild rodents can help researchers understand the links between specific genome changes and the ability to adapt to pressures in the environment, as well as better understand the evolution of rodents.

## Introduction

The study and characterization of genomic diversity of non-model organisms is complicated by limitations in knowledge and genomic resources available [1]. By contrast, researchers studying model organisms benefit from the advantage of working with species that have sequenced and annotated genomes, and high throughput platforms to survey genetic diversity at low cost. There is a lack of genomic sequence information available for non-model species, and a deficit of tools to assay genomic diversity in understudied organisms [2–4]. There is a need for custom tools to survey genomic diversity in non-model organisms, but the creation of these tools can be time consuming and expensive. There is an opportunity to explore existing technologies designed for model organisms and test the applicability of these tools in non-model species.

Genotyping arrays are convenient tools that obtain large amounts of genetic diversity information in a single assay at low cost [5]. Genotyping arrays are designed to capture a large swath of diversity within a species, but the technology is typically tailored to the model species of interest. Hybridization of microarray oligos targeted to unique locations in test DNA of the organism of interest provides a picture of the genomic landscape of that sample [6]. Single nucleotide polymorphisms (SNPs) are single base pair genome variations found in at least one percent of individuals in a population, and are an informative type of genomic diversity that is captured by genotyping arrays [6,7]. SNPs are found in abundance throughout the genome, and this variation can be used as a metric of genomic diversity when comparing different individuals in a population, or different species of interest [8].

There is a precedent for exploring the possibility of applying existing genotyping array technologies to related, non-model species. The majority of research examining the applicability of existing mammalian genotyping arrays in cross-species analyses focus on applying array technologies designed for agricultural and domestic breeding purposes to related species [2–4,9–14]. Researchers using a bovine genotyping array were able to identify a panel of over 100 candidate SNPs conserved within two species of wild oryx, despite a 23 million year divergence time between oryx and modern cows [2]. Other researchers have applied domestic arrays to non-model organisms that diverged from the model species millions of years ago to identify SNPs associated with an ideal physical trait that would inform breeding strategies [4], or to identify sexually selected traits that are associated with the fitness of a non-model organism [11].

Looking at the research performed in the field of cross-species genotyping array use, we identify three metrics of success for the application of existing genotyping arrays to non-model species. The first metric of success for applicability of genotyping arrays cross-species is the identification of a panel of candidate SNPs that may be conserved between the model and non-model organisms. This panel of SNPs represents variation that can be successfully genotyped in the non-model organism of interest. While one metric of success for genotyping array use is the number of loci or positions in the genome that can be accurately genotyped, the ability to detect heterozygous loci is the second metric. Heterozygous loci, or positions in the genome in which both the major and minor alleles in a population can be genotyped, are key when surveying diversity in populations [2,3,15,16]. The third and final metric of success we have identified is the ability to validate the candidate panel of SNPs and heterozygous loci either through *in silico* methods for non-model species with some sequence information available, or by testing for the candidate SNPs in populations using alternative experimental methods.

Genotyping arrays have demonstrated utility in identifying polymorphic SNPs, or sites of variability within non-model organisms, which is an important goal for conservation studies of endangered species, and molecular ecology [2,3,17]. In one particular study, researchers Hoffman et al. (2013) applied a Canine HD Beadchip genotyping array to a population of Antarctic fur seals, despite a 44 million year divergence time between the species of seal and dogs [3]. Using the Canine HD Beadchip which queries over 173,000 SNP loci in dogs, the researchers were able to identify a panel of 173 polymorphic SNP loci that were conserved between the Antarctic fur seals and dogs [3]. A subset of the loci genotyped were validated *in silico* using available transcriptomic data. Gene ontology analysis of shared loci between dogs and seals showed that the panel of loci were involved in energy metabolism, suggesting the genomic markers conserved between dogs and seals were a part of a highly conserved functional pathway.

The identification of SNPs in non-model species can be used as markers of rapid evolution between populations [18], and a genotyping array would allow researchers to identify large lists of candidate SNP loci in a single application. The characterization of SNPs across the genomes of wild organisms is of keen interest to population geneticists as molecular markers for comparative studies [19]. Cross-species genotyping can provide information regarding variants that are involved in sexual selection [11], and variants tied to a phenotype of interest, which can inform breeding strategies [4]. A study by More et al. (2019) tested the utility of a Bovine SNP50 array on the alpaca species *Vicugno pacos* which is bred domestically for its hair fiber that is economically valued [4]. The cross-species application of the Bovine SNP50 array allowed researchers to identify a panel of 400 polymorphic SNPs in the alpaca, and they were able to map 209 SNPs to alpaca gene sequence information that was available [4]. This study helped identify a number of SNP markers with utility cross-species that is currently needed to help guide breeding practices for the alpaca species that will maximize high-quality hair fiber yield in the future. This study also highlights the need for the development of genomic tools capable of genotyping non-model species of interest.

There are a number of different genotyping arrays that have been applied cross-species to non-model organisms, but there has been little research focusing on cross-species genotyping in mice and other rodents. Mice are a peculiarity in that most genomic tools are designed for classical, inbred mice used in research, but mice and related rodents can be found all across the world. There is a need for a tool that can survey diversity in non-model mice and other rodents. Wild rodents represent unique research opportunities because of the unique selective pressures that are placed on them through human influence, and their ability to rapidly adapt to changing human environments [18,20]. For instance, deer mice from the genus Peromyscus make interesting candidates for non-model research as they can be found across North America and despite lacking fully sequenced and annotated reference genomes, they have been previously used as sentinels of environmental contaminants [21], and are becoming key organisms for evolutionary studies and molecular genetics [18,22,23]. While some genetic resources are available for deer mice and other rodents of interest, there remains a paucity of genomic information for these understudied species and few low-cost tools to assess genomic variation in a high-throughput manner.

The Mouse Diversity Genotyping Array (MDGA) is a tool designed to survey hundreds of thousands of SNP loci across the genome of the house mouse and was specifically created to maximize the amount of SNP diversity that can be identified within laboratory mouse strains and crosses [24]. After testing and the removal of poorly performing SNP probes, the MDGA was found to genotype 493,290 SNP loci within the genome of the house mouse [25]. The aim of our study is to explore the use of the MDGA for its utility as a cross-species genotyping tool. The MDGA was tested on 44 samples ranging in relatedness to the model house mouse, *Mus musculus*, that span different Genus, Family, and Orders of taxonomic classification (Table 1, S1 Table). The goal was to identify the three metrics of success that define MDGA cross-species utility in related organisms. This study represents an advance in the field of mammalian cross-species genotyping that will add to the paucity of genomic sequence and SNP information available for non-model mice and rodents (Fig 1). It was hypothesized that application of the MDGA to wild rodent DNA samples will help elucidate potential polymorphic loci, or the number of loci that can detect both the A and B allele in a population, and that can be used cross-species in non-model organisms.

**Table 1.**
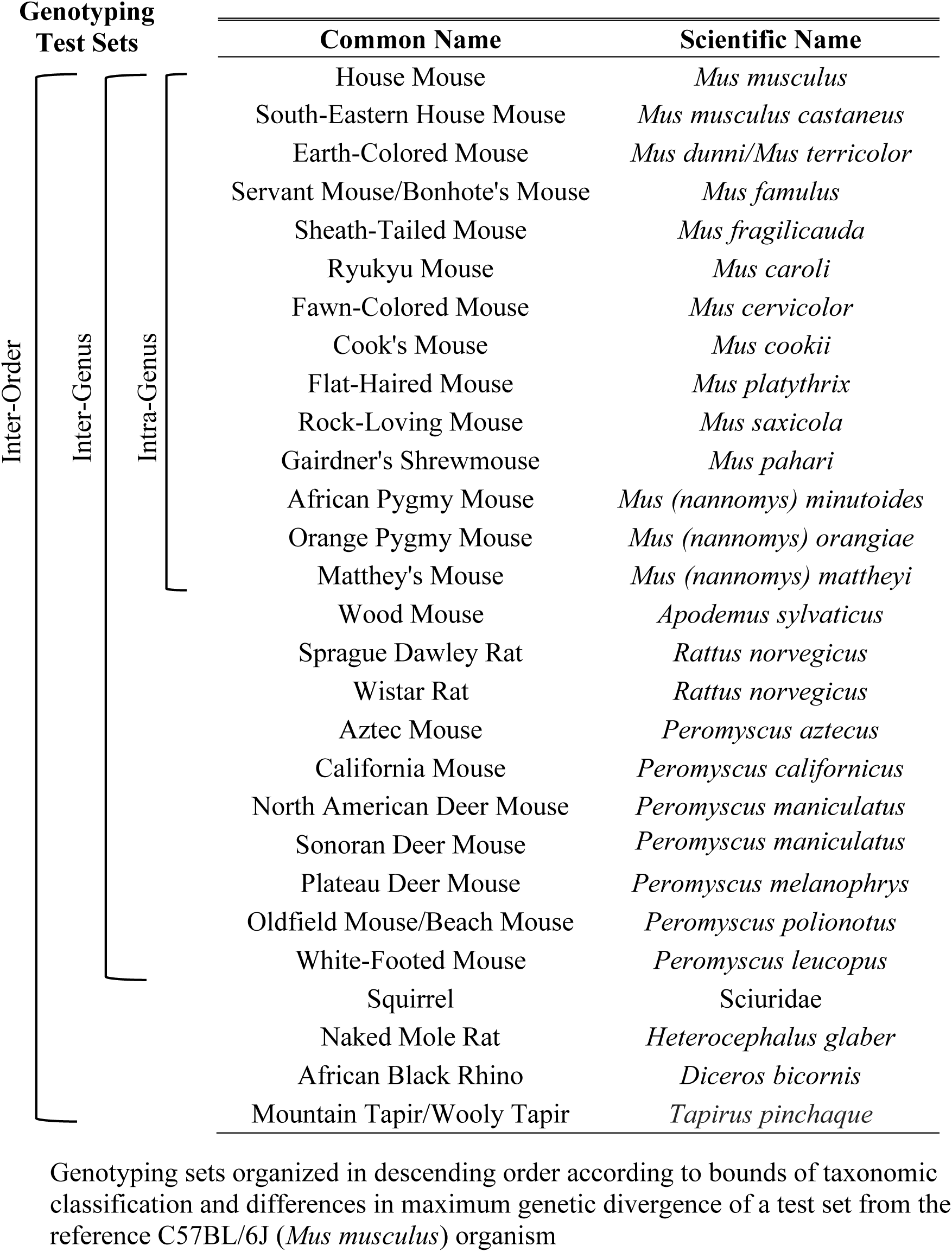
Genotyping sets of study

**Fig 1.**
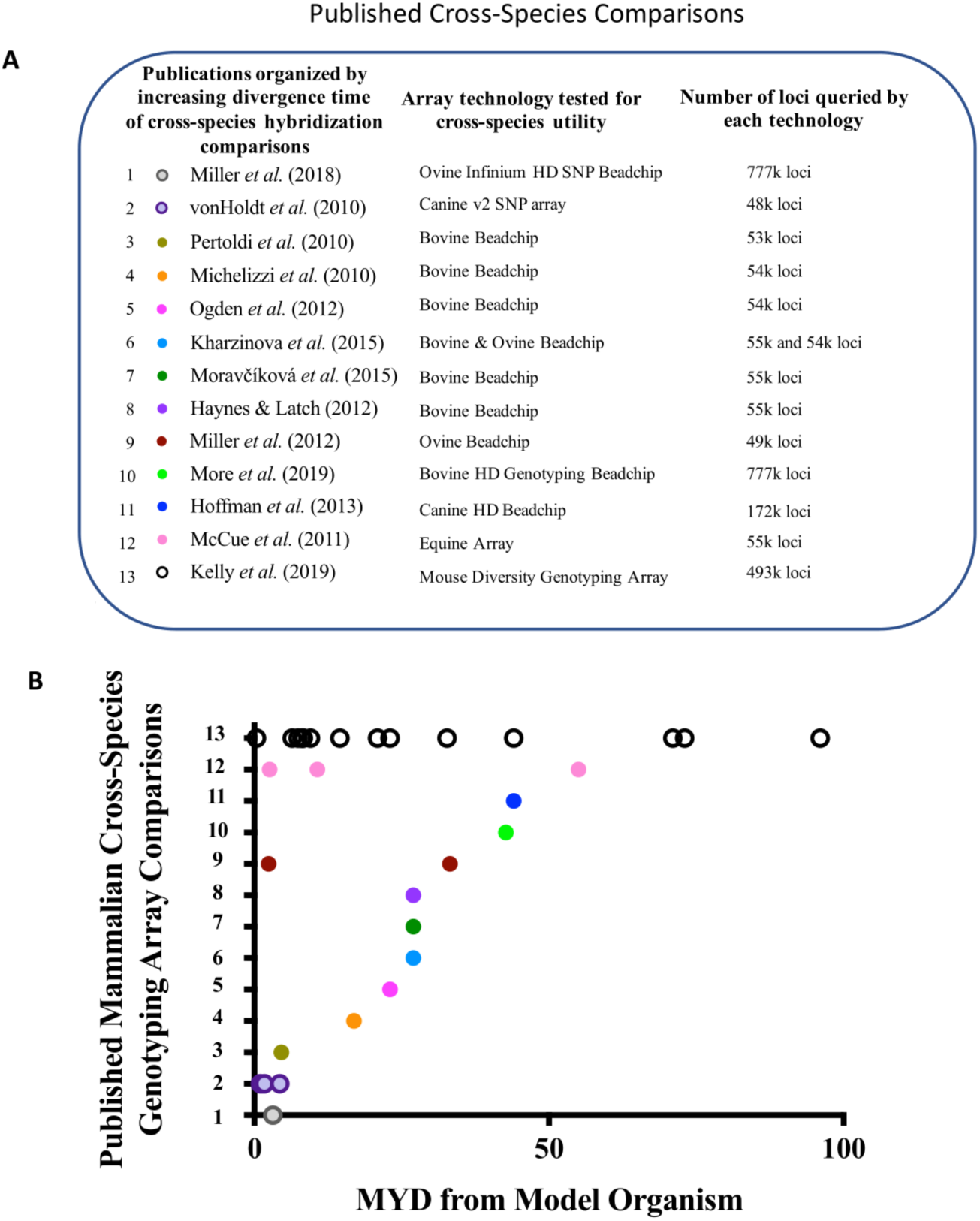
Summary of published research on mammalian cross-species genotyping using SNP genotyping microarrays. (A) Published research is organized in increasing order of genetic divergence in millions of years divergence (MYD) of non-model test samples from the model reference organism. Authors, publication year, genotyping microarray technology, and approximate number of loci queried (in thousands) are listed for each publication. (B) The sample of publications on mammalian cross-species array studies with the 13^th^ representing the contributions of this thesis to the cross-species genotyping array field.

## RESULTS

### Cross-species test sets exceed maximum genetic diversity of the training set

A training set of DNA samples from 114 classical, inbred laboratory mice was used in training the genotyping algorithm employed by Affymetrix Power Tools to provide accurate genotypes (S2 Table). Genetic distances reflect the relatedness between samples and were obtained from calculations of SNP distances derived from raw genotyping results. The maximum genetic distance of the training set is approximately 0.225 with respect to the reference C57BL/6J house mouse (Fig 2). The intra-genus test set of 27 species from the genus Mus has a maximum genetic distance value of 0.836 and is over three times larger than the maximum genetic distance of the reference set of 114 classical inbred mice (Fig 2). A case study of seven Peromyscus samples genotyped together has a maximum genetic distance of 0.941 from the house mouse, and far exceeds the diversity of the training set. Also, the maximum genetic distance of the inter-order test set (n=44, 96 MYD) is 0.938, and is over four times larger than the maximum genetic diversity represented in the training set (Fig 2). The training set used does not encapsulate the genetic diversity of the test sets.

**Fig 2.**
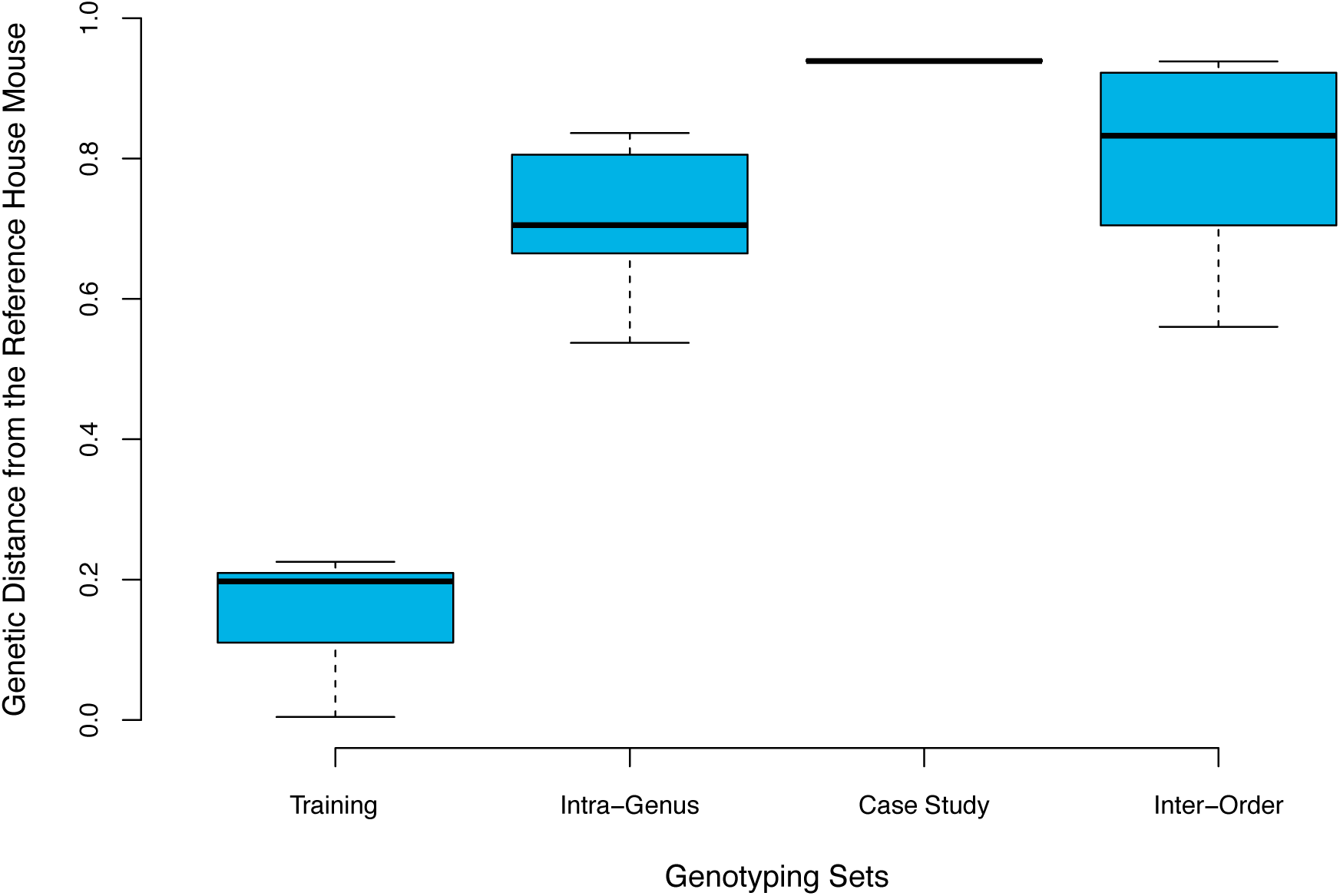
Genetic diversity of test sets exceeds maximum genetic diversity of training set. Boxplots representing the minimum, first quartile, median, third quartile, and maximum genetic distances for the training set (n=114), intra-genus test set (n=27), case study of Peromyscus (n=7), and inter-order test set (n=44). All genetic distances are with respect to the reference house mouse *Mus musculus*.

The samples of the inter-order test set are significantly different in genotypic composition and allelic frequency (P<0.0001; Fisher’s exact test with Monte Carlo simulation). The samples of the intra-genus test set of only Mus samples are also significantly different in genotypic and allelic frequency (P<0.0001). Two *R. norvegicus* samples were compared to one another as a control and the genotypic composition is not significantly different (p=0.0934). Differences in allelic composition between *R. norvegicus* samples are also not significant (p = 0.2232). The four *H. glaber* (naked mole rat) samples genotyped together are significantly different in the genotype composition (p<0.0001), but not allelic composition (p=0.0038).

### Underestimation of genetic diversity occurs when genotyping across multiple genera

For the inter-order genotyping set (n=44), a general decrease is observed in the percentage of loci genotyped as divergence time increases from *M. musculus* (r = −0.57; p-value<0.0001; Fig 3A). As divergence time increases from *M. musculus*, the number of ‘no calls’, or inability to determine a genotype at a locus, increases. A plateau in the percentage of loci genotyped is observed between 10-15 MYD for non-Mus samples from the inter-genus test set. Loci with heterozygous genotypes were of particular interest, as those loci have the potential to identify both the major and minor alleles in a population (polymorphic loci). The percentage of loci that had a heterozygous genotype increases as divergence time from the house mouse increases (Fig 3B). There is a positive correlation between increasing percent heterozygosity and the known divergence times from the house mouse (r = 0.67; p-value<0.0001). Similar to the percentage of loci genotyped, a plateau in percent heterozygosity is also observed to begin between 10-15 million years divergence from *M. musculus* (Fig 3B).

**Fig 3.**
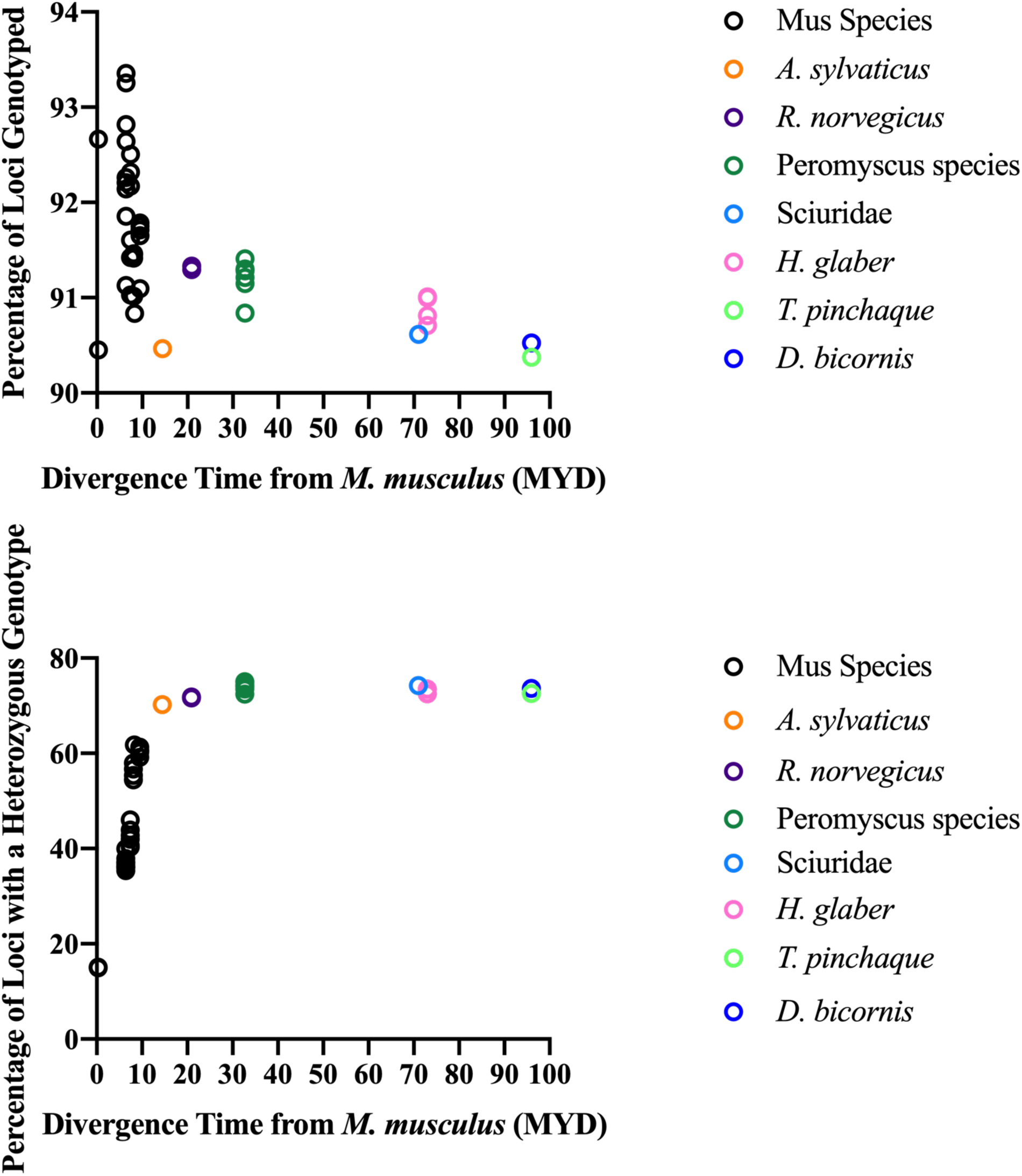
Underestimation of genetic diversity for highly diverged species in cross-species genotyping. (A) The percentage of loci genotyped from the inter-order test set (n=44). (B) The percentage of loci from the inter-order test set with a heterozygous genotype call. MYD = Millions of years divergence, with respect to the reference *Mus musculus*.

### MDGA captures the genetic diversity of wild samples from the genus Mus

As seen in the inter-order test set, there is a general decrease in the percentage of loci that were genotyped in samples of the intra-genus test set (Fig 4A). There is a negative correlation between the percentage of loci genotyped and the known divergence times from *M. musculus* (r = −0.76; p<0.0001). In the intra-genus test set, heterozygosity increases as divergence time increases (Fig 4B). The increase in percent heterozygosity of Mus samples is positively correlated with an increase in divergence times (r = 0.93; p-value<0.0001). There is no plateau or obvious underestimate of genetic diversity for samples in the intra-genus test set.

**Fig 4.**
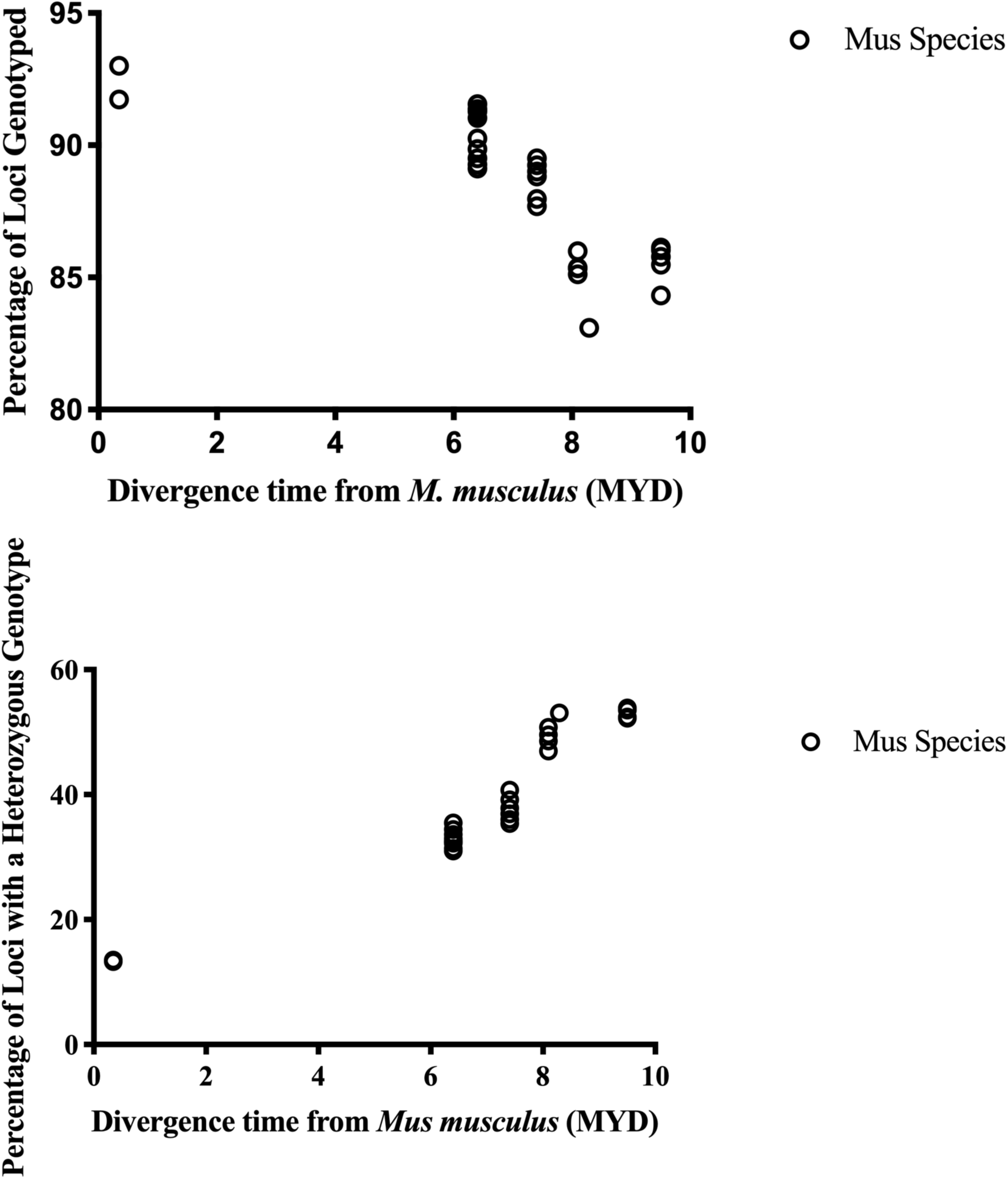
Genetic diversity of wild Mus species. (A) The percentage of loci genotyped from the intra-genus test set (n=27). (B) The percentage of loci from the intra-genus test set with a heterozygous genotype call. MYD = Millions of years divergence, with respect to the reference *Mus musculus*.

A tree of relatedness derived from SNP-based genetic distance values differentiates Mus samples of the intra-genus test set from one another at a species level (Fig 5). Enough genetic diversity is captured using the MDGA to reflect the known taxonomic relationships between the intra-genus samples at a species level. At 9.5 MYD, the pygmy mouse subspecies *M. n. minutoides* is grouped with the subspecies *M. n. orangiae* and not the replicate data file of the same species.

**Fig 5.**
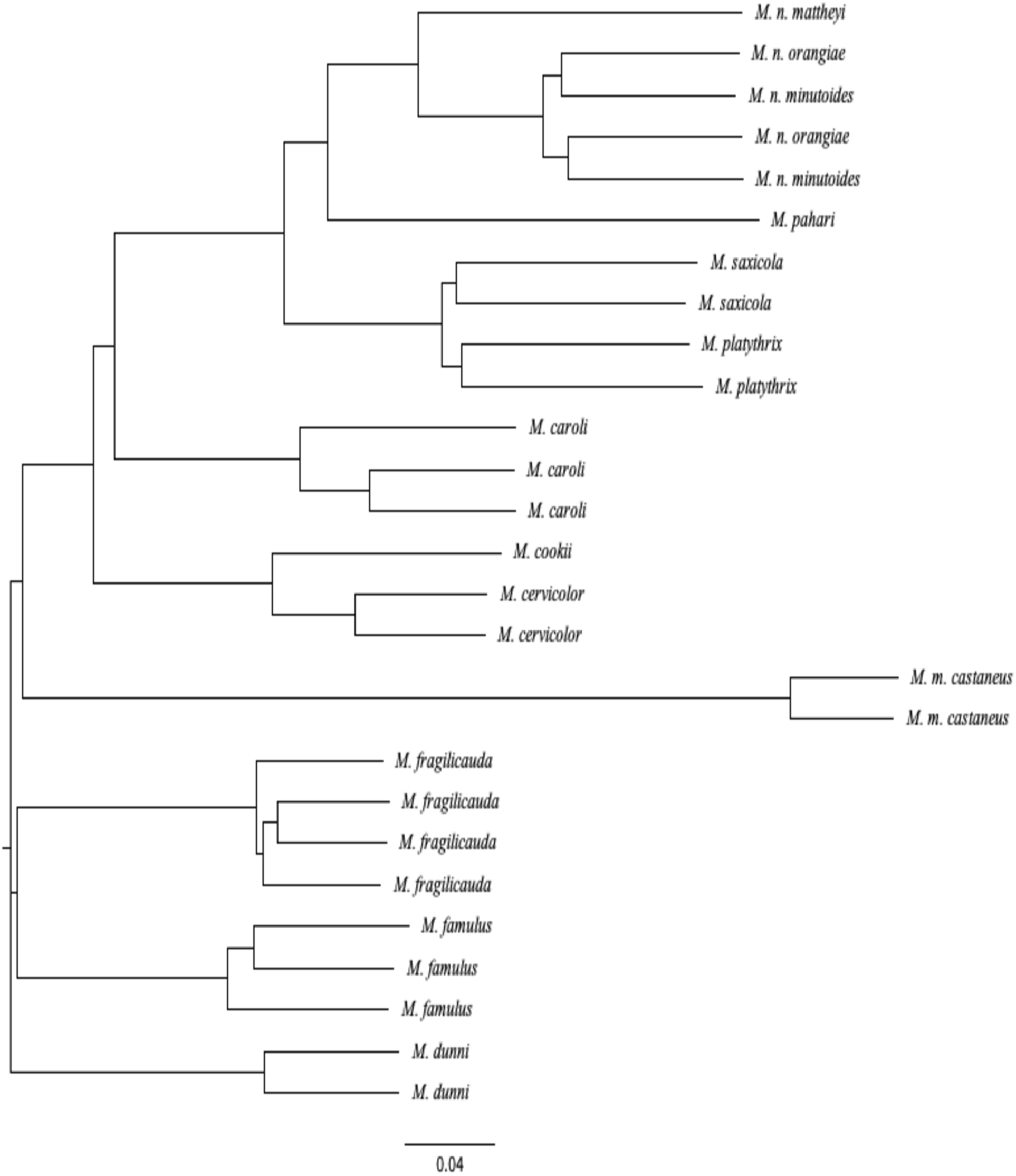
SNP distance-based tree of genetic relatedness reflects known taxonomic relationships between Mus species. SNP distance-based tree of genetic relatedness of samples from the intra-genus test set (n = 27). At 9.5 MYD a pygmy mouse subspecies *M. n. orangiae* has SNP-based genetic distances that reflect greater genetic similarity to another pygmy mouse subspecies *M. n. minutoides* than the replicate MDGA data file of the same *M. n. orangiae* sample. MYD = Millions of years divergence, with respect to the reference *Mus musculus*.

### Peromyscus case study

Seven Peromyscus species were genotyped together as a case study to determine if the MDGA could provide useful results that reflect known biological diversity for a number of species of a different genus from Mus. Of the Peromyscus samples queried, approximately 52% of loci queried by the array produce a genotype (Table 2). There are 159,797 loci genotyped across all seven samples (32% of loci queried by the array) despite a 32.7 million-year divergence time from *M. musculus*. SNP-based genetic distances of Peromyscus species were utilized to produce trees of genetic relatedness that reflect the known divergence times of these species (Fig 6). Top KEGG pathway annotations of the genotyped loci in Peromyscus samples are associated with neurological signaling (Table 3).

**Table 2.** Percentage of loci genotyped and percent heterozygosity in a Peromyscus case study (n=7)

| MDGA Data (CEL) File | Sample Scientific Name | Loci Genotyped (%) | Heterozygosity (%) |
| --- | --- | --- | --- |
| SNP_mDIV_B2-660_102109.CEL | *P. melanophrys* | 51.31 | 34.83 |
| SNP_mDIV_B1-659_102109.CEL | *P. aztecus* | 52.03 | 36.02 |
| SNP_mDIV_B3-661_102109.CEL | *P. californicus* | 52.13 | 36.27 |
| SNP_mDIV_B4-662_102109.CEL | *P. m. sonoriensis* | 52.26 | 35.95 |
| SNP_mDIV_B5-663_102109.CEL | *P. m. bairdii* | 52.27 | 36.71 |
| SNP_mDIV_B6-664_102109.CEL | *P. polionotus* | 52.57 | 37.02 |
| SNP_mDIV_B8-666_102109.CEL | *P. leucopus* | 52.62 | 36.55 |

**Table 3.**
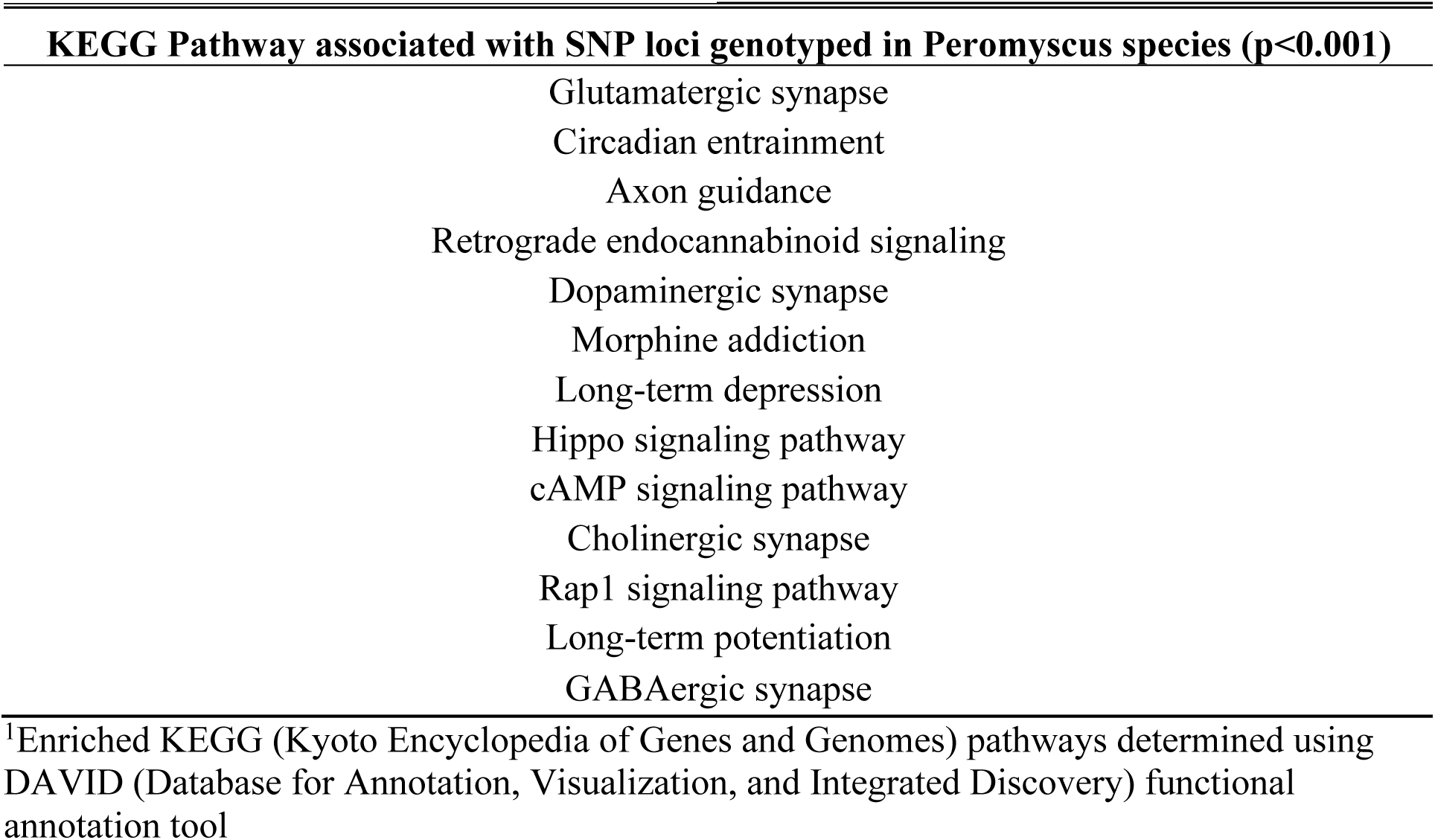
Top KEGG^1^ (Kyoto Encyclopedia of Genes and Genomes) pathways determined using the DAVID functional annotation tool

**Fig 6.**
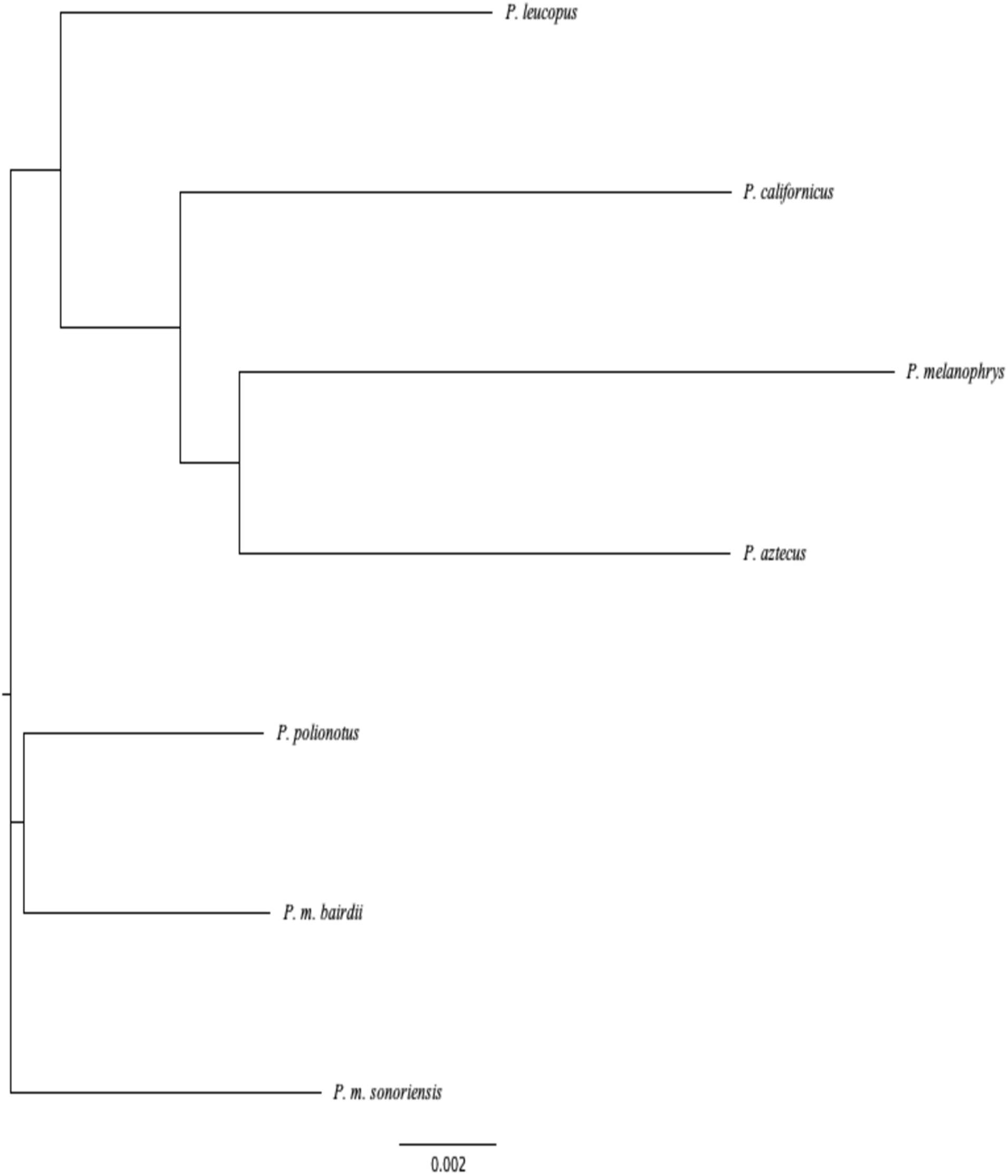
SNP distance-based tree of genetic relatedness reflects known taxonomic relationships between Peromyscus species. Pairwise SNP distance-based tree of genetic relatedness of samples from the intra-genus test set of Peromyscus species (n=7).

### *In silico* cross-validation of potential polymorphic loci

*P. maniculatus* was examined given that there is a partial genome sequence available online for *in silico* search of unique and perfect 25 nt MDGA probe target sequence matches. There are 226,265 loci on the MDGA genotyped (∼52%) for both *P. maniculatus bairdii* and *P. maniculatus sonoriensis* within this study. Of the loci that were genotyped, there are 143,971 loci that were genotyped as heterozygous in both *P. maniculatus* samples (Table 4). Heterozygous loci represent potential polymorphic loci that can query both the common and uncommon allele in a population. There are 6,075 MDGA probe sequences that perfectly match a unique position within the *P. maniculatus* genome, and 481 of the *in silico* sequence matches are associated with heterozygous loci.

**Table 4.** In silico validation of potential polymorphic loci conserved cross-species

| Common Name | Scientific Name | MYD from *M. musculus* | Unique in silico matches of alleles queried | Number of heterozygous loci in all samples | Number of candidate polymorphic loci |
| --- | --- | --- | --- | --- | --- |
| Ryukyu Mouse | *Mus caroli* | 7.41 | 303,680 | 147,452 | 9,413 |
| Gairdner's Shrewmouse | *Mus pahari* | 8.29 | 152,971 | 251,902 | 9,341 |
| Rat | *Rattus norvegicus* | 20.9 | 61,372 | 85,926 | 1,019 |
| Deer Mouse | *Peromyscus maniculatus* | 32.7 | 6,075 | 143,971 | 481 |
| Naked Mole Rat | *Heterocephalus glaber* | 73 | 1,179 | 91,324 | 52 |

MYD=Millions of Years Divergence
Candidate Polymorphic Loci are the number of loci identified that had a heterozygous genotype call for the samples using the Mouse Diversity Genotyping Array Data that could also be mapped to the available genomic sequences for these organisms.

An average of 382,968 loci were genotyped between three available *M. caroli* CEL files using the MDGA, and there are 303,680 unique theoretical matches to the *M. caroli* genome determined through an *in silico* search using E-MEM (Table 4). A shrew mouse (*M. pahari*) applied to the array has 411,514 loci that were genotyped experimentally using the MDGA. Theoretically, there are 152,971 unique sequences from the MDGA that are present in the shrew mouse only once (Table 4). The pathways associated with genotyped loci in *M. musculus, M. caroli*, and *M. pahari* that are shared between these three species are primarily signaling pathways and pathways involved in maintaining the structural integrity of a cell, such as focal adhesion and adherens junction (Table 5). The Sprague Dawley rat (*R. norvegicus*) has a fully sequenced and annotated genome available online. There are 170,156 loci that were genotyped experimentally in both *R. norvegicus* samples using the MDGA. Using the E-MEM *in silico* program, 61,372 sequences were determined to be theoretically present within the genome (Table 4).

**Table 5.**
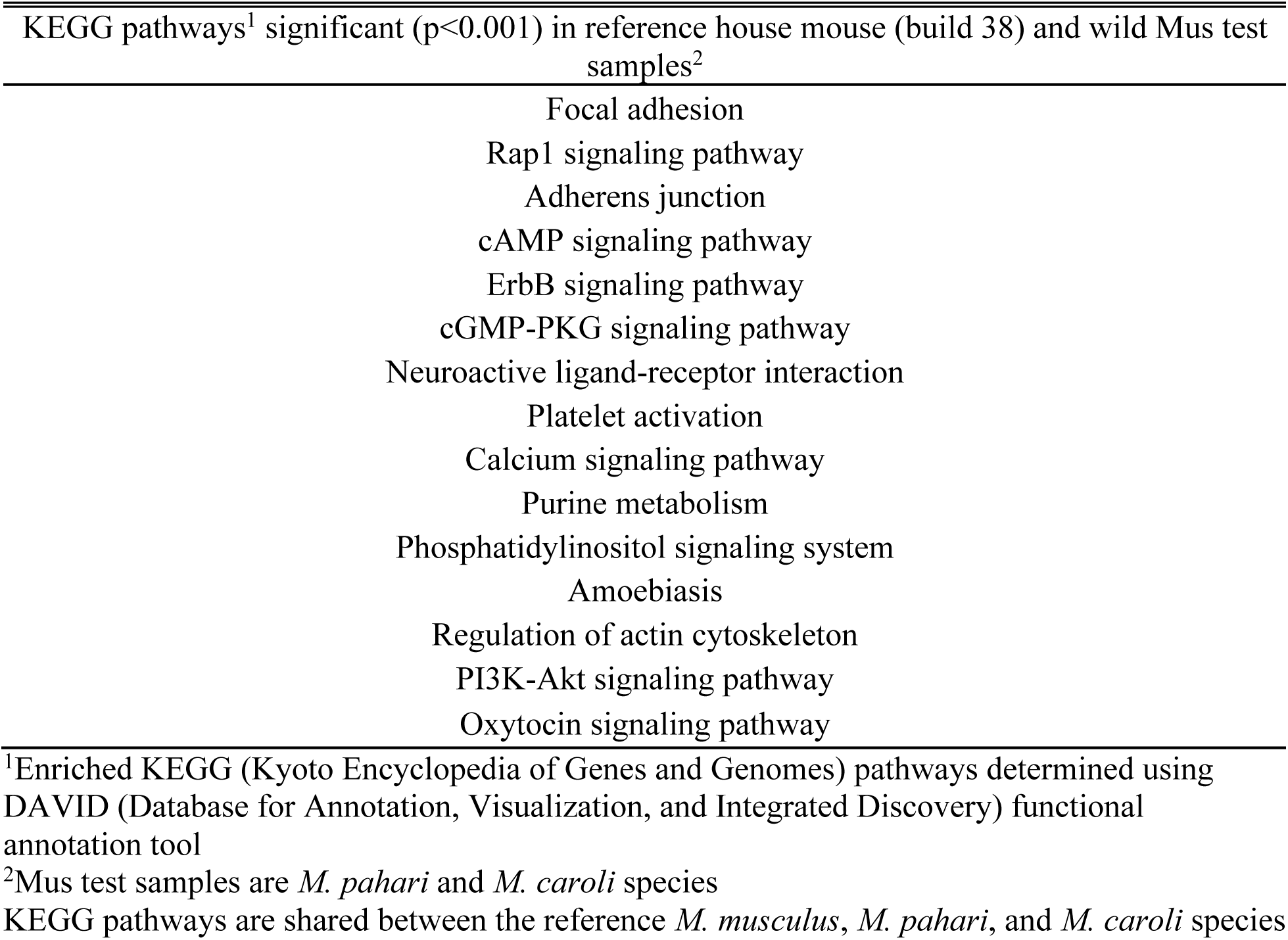
Top KEGG pathways enriched for house mouse gene annotations with genotype assignments across wild Mus species

| KEGG pathways<sup>1</sup> significant (p<0.001) in reference house mouse (build 38) and wild Mus test samples<sup>2</sup> |  |
| --- | --- |
|  | Focal adhesion |
|  | Rap1 signaling pathway |
|  | Adherens junction |
|  | cAMP signaling pathway |
|  | ErbB signaling pathway |
|  | cGMP-PKG signaling pathway |
|  | Neuroactive ligand-receptor interaction |
|  | Platelet activation |
|  | Calcium signaling pathway |
|  | Purine metabolism |
|  | Phosphatidylinositol signaling system |
|  | Amoebiasis |
|  | Regulation of actin cytoskeleton |
|  | PI3K-Akt signaling pathway |
|  | Oxytocin signaling pathway |

<sup>1</sup>Enriched KEGG (Kyoto Encyclopedia of Genes and Genomes) pathways determined using DAVID (Database for Annotation, Visualization, and Integrated Discovery) functional annotation tool
<sup>2</sup>Mus test samples are *M. pahari* and *M. caroli* species
KEGG pathways are shared between the reference *M. musculus*, *M. pahari*, and *M. caroli* species

Special attention was given to potential polymorphic loci that were genotyped as heterozygous in samples using the MDGA and could be cross-validated as being present in the genome using an *in-silico* search of publicly available genome sequences. There is a trend of there being more heterozygous loci genotyped using the MDGA than the number of those loci that can be cross validated as present in the publicly available genome sequence (Table 4). There are 147,452 heterozygous loci genotyped in all three *M. caroli* samples, and 9,413 of these loci were validated as present in the publicly available genome sequence (Table 4). There are 9,341 of the 147,452 heterozygous loci genotyped in a *M. pahari* sample that were cross validated as potential polymorphic SNP loci (Table 4). In two *R. norvegicus* samples, there are 85,926 loci that were genotyped empirically using the MDGA, and 1,019 loci that were cross-validated using an *in-silico* genome sequence search (Table 4).

## Discussion

Specialized genotyping arrays have been successfully applied cross-species to closely related organisms in previous research [2–4,9–13,16,32–34]. Here we present evidence that the MDGA can be applied to wild rodents to produce SNP genotyping results that reflect the known taxonomic relationships between test samples and the reference house mouse. The identification of polymorphic SNPs within non-model organisms is of great interest, as these genetic markers can be used to assay diversity in wild populations in studies of population genetics [2–4,9,16,18,34,35]. Panels of candidate polymorphic SNPs have been identified for wild species of the genus Mus and Peromyscus. This study is a first step in contributing where there is a paucity of information available for non-model rodent species.

Outside of the genus Mus, the plateau in SNP loci genotyped and the percentage of heterozygous loci is attributed to off-target mutations that hinder DNA hybridization to array probe sequences. When DNA hybridizes to a probe on the MDGA, the hybridization does not have to be a perfect 25 nt match, where incomplete hybridization of the sample DNA to the probe is enough to result in a genotype assignment [36]. Determination of the divergence time from *M. musculus* at which genetic diversity is underestimated is limited by the samples available for use in this study. A greater number of species genotyped using the MDGA that have a divergence time between 10-15 MYD from the house mouse would be beneficial in identifying situations where underestimations of genetic diversity occur. Miller et al. (2012), found previously that applying the Bovine, Ovine, and Equine SNP50 Beadchip arrays cross-species resulted in a linear decrease in genotyped loci as the millions of years of divergence from the model species increased [10]. Previous studies that have examined the utility of the cross-species application of commercially available genotyping array technology have identified trends of decreasing ability to genotype loci as divergence time from the model organism increases as well [8,9]. This study is unique as it tests the array technology on a wide range of species spanning multiple millions of years divergence from the reference house mouse.

Previous research has determined potentially conserved sequences between model organisms and the wild species of interest through application of commercial arrays to test samples [2,4]. This study of the MDGA cross-validates genotyped loci in rodent samples with an *in silico* analysis of available genomic sequences for wild species. The heterozygous SNP variation in rodent samples of this study cross-validated through *in silico* analyses represents candidate polymorphic SNPs that can be tested for conservation in populations of wild species of Mus and Peromyscus. To be truly considered a polymorphic SNP conserved cross-species, the variation must be validated in wild populations with the alternate, or minor allele present in at least 1% of the population.

A major difficulty in cross-species genotyping using array data is the assembly of appropriate test sets that would allow for accurate genotyping. Previous research has demonstrated that the genotyping algorithm recommended by Affymetrix, BRLMM-P, is sensitive to the composition of the samples included in a test set [37,38]. Samples in a test set that are more similar to one another genetically will produce fewer false genotyping results [38]. The number of loci genotyped can become inflated if the samples in the test set are too genetically different, as was seen when samples of different orders of classification were genotyped together in the inter-order test set. The greater genetic homogeneity of only Mus samples in the intra-genus test set produced genotyping results that matched what was expected of the species based on divergence times. The linear decrease in loci genotyped in Mus samples as divergence time increased reflected previous cross-species findings [10]. Recommendations for the construction of a test set of samples for an experiment utilizing the MDGA cross-species would be dependent upon the hypothesis tested. A large number of samples are needed to establish whether the minor allele of a SNP is present in populations of non-model species for at least 1% of the population [7,39]. Technical replicates should be included to assess the quality of DNA hybridization to array probes for a particular species. Optimization of hybridization conditions should be made to reduce differences in array hybridization intensities and the resulting differences in genotype assignments between technical replicates.

The use of a training set that has sufficient genetic diversity to encompass that of the experimental test sets can assist in producing accurate genotyping of samples [40,41]. The training set of 114 classical inbred strains of mice used in this study does not encompass the high relative genetic diversity of the sample sets of this cross-species study. A training set optimized for cross-species genotyping would be composed of members of the same species as the test set and would be validated using another method such as sequencing. Inclusion of male and female samples would ensure more accurate genotype assignments on the X chromosome, as hemizygous males are assigned a diploid homozygous genotype [42]. Analyzing SNPs on the X chromosome separately from autosomal SNPs and separating male and female samples would aid in fewer false genotype assignments.

In comparing the research knowledge gained through this study using the deer mouse (*P. maniculatus*) to the knowledge obtained from the study of Antarctic fur seals by Hoffman et al. (2013), similar metrics of utility were obtained through cross-species genotyping (Table 6). Given that the mouse array targets over two times the number of positions than the canine array targets, there is a much larger number of loci that can be genotyped in the deer mouse than the Antarctic fur seal. Future studies will focus on validating a panel of SNPs that are polymorphic in deer mouse populations. Pathway analyses are limited by the information assayed by each technology and are with respect to the annotations of the model organisms. As new sequence information and genome annotations become available for the deer mouse, it will be interesting to see which SNP markers associated with conserved pathways will be found to be shared between the house mouse and the deer mouse. The deer mouse is an intriguing sentinel of environmental effects and a model for population studies that has a surprising lack of genomic information available [18,43]. Cross-species array use may be one technique to identify SNP diversity in these relevant species until genome sequencing prices become more affordable for non-model species. The use of a rat genotyping array in the future may be of use, as the deer mouse and rat share greater genetic synteny than with the mouse [44].

**Table 6.** Comparison of the *Hoffman et al. (2013)* model study with the current study.

| Hoffman et al. | Comparison | Kelly et al. |
| --- | --- | --- |
| Antarctic fur seal | Non-model species | Deer Mouse |
| CanFam2.0 | Reference genome for array | *Mus musculus* |
| 44 | MYD from model species | 32.7 |
| 173,662 | Loci queried by array | 493,290 |
| 33,324 | Loci genotyped | ~226,000 |
| 2 of 5 | Loci validated in silico | 3,195 |
| 173 | Polymorphic loci | 481 |
| 2 | Loci validated in a population | Future |
| Energy metabolism | Pathways shared between model and non-model species | Neurological signaling |

MYD = Million years divergence

There is a great potential for cross-species MDGA utility for wild Mus species in providing genomic markers for research in mouse population genetics and studies of rodent evolution. Genotype data generated from application of the MDGA captured enough genetic diversity to differentiate Mus samples at a species level. Further testing is required to determine if the MDGA can capture enough diversity to differentiate between subspecies. As in the case of the deer mouse, wild Mus species represent an untapped wealth of genomic information that would benefit researchers of environmental mutagens, evolution, and population genetics. With newer mouse array technologies becoming available that have greater capacity for high-throughput analysis, novel polymorphic SNPs in non-model rodents can be identified through a low-cost and efficient manner.

Utilizing the Mouse Diversity Genotyping Array for cross-species genotyping represents a first step towards development of a tool that can rapidly identify SNP variation in wild rodent species. A panel of candidate SNPs on the MDGA have been identified for use with wild mouse species and was cross validated using an *in silico* genome search. Future work may address the validation of this candidate cross-species panel in wild populations. This research highlights the need for greater genomic resources for wild rodents and demonstrates the potential of the MDGA as a high-throughput genotyping tool for non-model organisms. The development of novel tools specialized for non-model species opens up previously inaccessible avenues of research. Next-generation sequencing technologies are often not accessible and too costly for a majority of researchers with population-based research questions that require rapid, high-throughput genome wide analysis of variation. Until the price of sequencing and the complexity of assembling new reference genome assemblies is reduced, the adaptation of existing genomic tools for use with closely related species is one method researchers can use to combat the genomic disparity between studying model and non-model species.

## MATERIALS AND METHODS

### Cross-species samples

Forty publicly available MDGA raw data (CEL) files were downloaded from the Center for Genome Dynamics at the Jackson Laboratory (2012, The Jackson Laboratory; ftp.jax.org/petrs/MDA/). Four MDGA CEL files of *H. glaber* DNA cross-hybridization to the MDGA were generated in-house. The forty-four samples consist of twenty-seven Mus CEL files, two Rattus CEL files, seven Peromyscus CEL files, one Apodemus CEL file, and CEL files representing more highly diverged species including a squirrel, four naked mole rats, a tapir, and an African Black Rhino (S3 Table). CEL file raw array intensity images were analyzed for quality and abnormalities in array images were noted. Two CEL files (S1 Fig) were noted for having an abnormal spot with uneven DNA hybridization to the array. Due to the redundancy of probes on the MDGA, it was determined that abnormal CEL file images still had sufficient genomic coverage to be used for further analysis and were not removed from the study.

### SNP genotyping

Samples were genotyped using the protocol outlined by Locke et al. [25]. Affymetrix Power Tools was used to generate genotype calls of AA, AB, BB, or No Call (numerical representations 0, 1, 2, −1, respectively) using the BRLMM-P algorithm for 493,290 SNPs [25] (Affymetrix Power Tools (APT) Release 1.16.0). A training set of 114 classical laboratory mouse CEL files obtained from a set of 351 mice utilized by Didion et al. (2012) was used in conjunction with BRLMM-P to train the algorithm in accurate assignment of genotypes [26]. The samples were organized into three test sets that were genotyped separately from one another. The first genotyping set (known as the inter-order test set) consists of all 44 CEL files representing species spanning different orders of classification and a maximum divergence time of 96 million years of divergence (MYD) from the reference house mouse, *Mus musculus* (Table 1). The second test set (the intra-genus test set) is composed of the 27 samples from the genus Mus and has a maximum divergence of 9.5 MYD from the house mouse (Table 1). The third test set (Peromyscus case study test set) was composed of seven deer mouse species from the genus Peromyscus that have 32.7 MYD from the house mouse (Table 1). The genotyping results obtained were analyzed and compared to reference genotyping data from *Mus musculus*. The reference *Mus musculus* data was obtained by averaging the genotyping results from 8 *Mus musculus* samples (percentage of loci genotyped > 99%).

### Estimation of divergence times

The estimated divergence time of each species from the reference house mouse was obtained using an evolutionary timetree of life (http://www.timetree.org/) [27] with a few exceptions. The estimated divergence of the subspecies *M. m. castaneus* was determined through previous work by Geraldes et al. (2012) [28], and the evolutionary divergence time of the pygmy mouse species from the house mouse was determined by Kouassi et al. (2008) [29].

### Statistical analyses

A Fisher’s exact test was utilized to assess the extent of genetic differences between samples genotyped together. A nonparametric, unordered, Fisher-Freeman-Halton exact test (Monte Carlo simulation) was performed using the StatXact statistical analysis software package (CYTEL Software, Cambridge, MA). Pearson’s r was used in tests of significance of correlations between the genotyping results of the test set samples using Graphpad Prism 8 software.

### Genetic distance calculations

Pairwise comparison of SNP genotypes between species in the inter-order test set was utilized to create SNP-based distance matrices using R. The distance matrix values used to create phenograms (SNP trees) were generated using an in-house R script courtesy of Marjorie E. Osbourne Locke. The in-house script utilized the ‘bionj’ R package to create a tree of genetic relatedness using the neighbour-joining method [30]. The resulting trees were modified using Figtree (v1.4.3) software. Pairwise genetic distances were computed by dividing the total number of genotypic differences between two samples by the total number of loci queried by the MDGA, where 493,290 total loci were used in this study [25]. The values in the distance matrix are a numerical representation of the amount of genetic diversity between test species analyzed and the reference house mouse. A genetic distance value of zero indicates the species are genetically the same at the loci queried, and a value of one indicates the species compared are completely genetically dissimilar from one another at the loci queried. The estimated evolutionary relationships seen in the SNP trees generated were compared to the divergence times of test samples from the reference house mouse provided in literature and the Timetree database [27–29].

### *In silico* validation of MDGA loci genotyped cross-species and pathway analysis

*In silico* validation of loci genotyped from MDGA data was performed using the program E-MEM (efficient computation of maximal exact matches for very large genomes) designed by Khiste and Ilie (2015) [31]. The publicly available genomes of rodents were searched for the unique presence of MDGA probe sequences. E-MEM was employed to search a publicly available genome of wild rodents available on NCBI (S4 Table) for perfect 25 nt MDGA SNP probe target sequences that have only one genomic match (ftp.ncbi.nlm.nih.gov/genomes/). Unique MDGA matches discovered via E-MEM were identified and then compared with the list of heterozygous loci genotyped using the MDGA. Ensembl gene IDs associated with candidate loci genotyped were analyzed using the Database for Annotation, Visualization, and Integrated Discovery (DAVID).

## Acknowledgements

Tail-tissue samples of four *Heterocephalus glaber* individuals were given to the Hill Laboratory by Dr. Melissa Holmes (Associate Professor at the University of Toronto, Mississauga Campus). CEL files of the rhino and tapir samples were donated by Karen Svenson from the Jackson Laboratory. Permission to use squirrel CEL file data was given to the Hill Laboratory by Dr. Fernando Pardo Manuel de Villena from the University of North Carolina at Chapel Hill. Assistance with E-MEM code was provided by Dr. Lucien Ilie (University of Western Ontario, London) and his student Qin Dong.

## Supporting Information

**S1 Fig.**
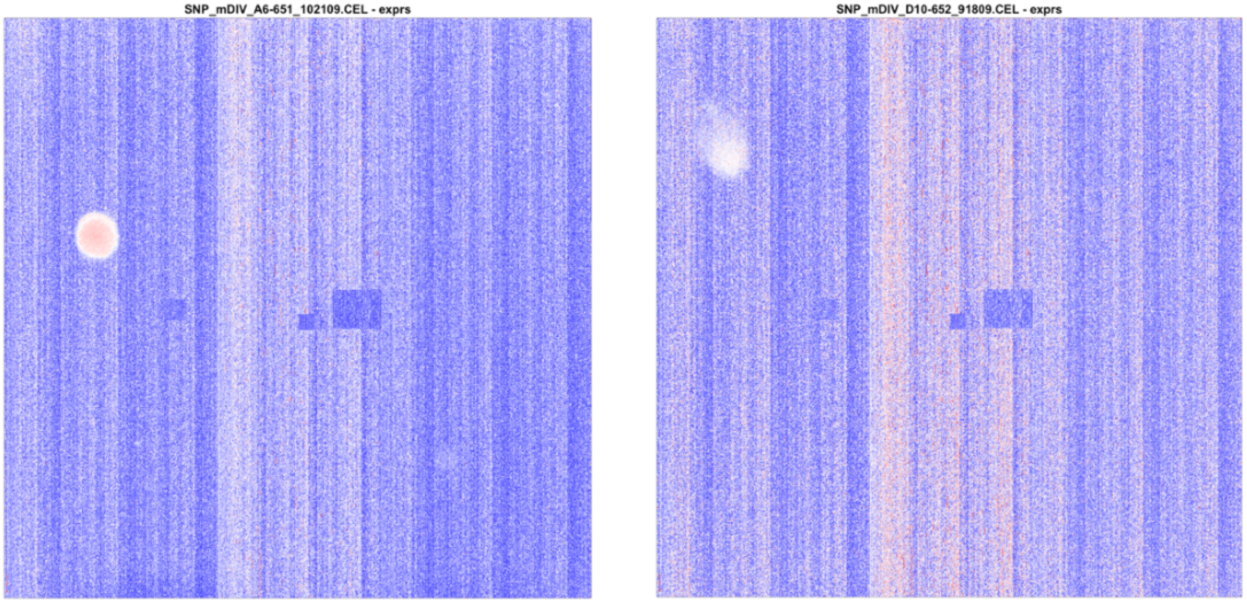
Abnormalities in two MDGA raw intensity CEL file images. CEL file raw array intensity images were analyzed for quality control purposes and abnormalities in array images were noted for two CEL files. The two samples were not removed from analysis.

**S1 Table.**
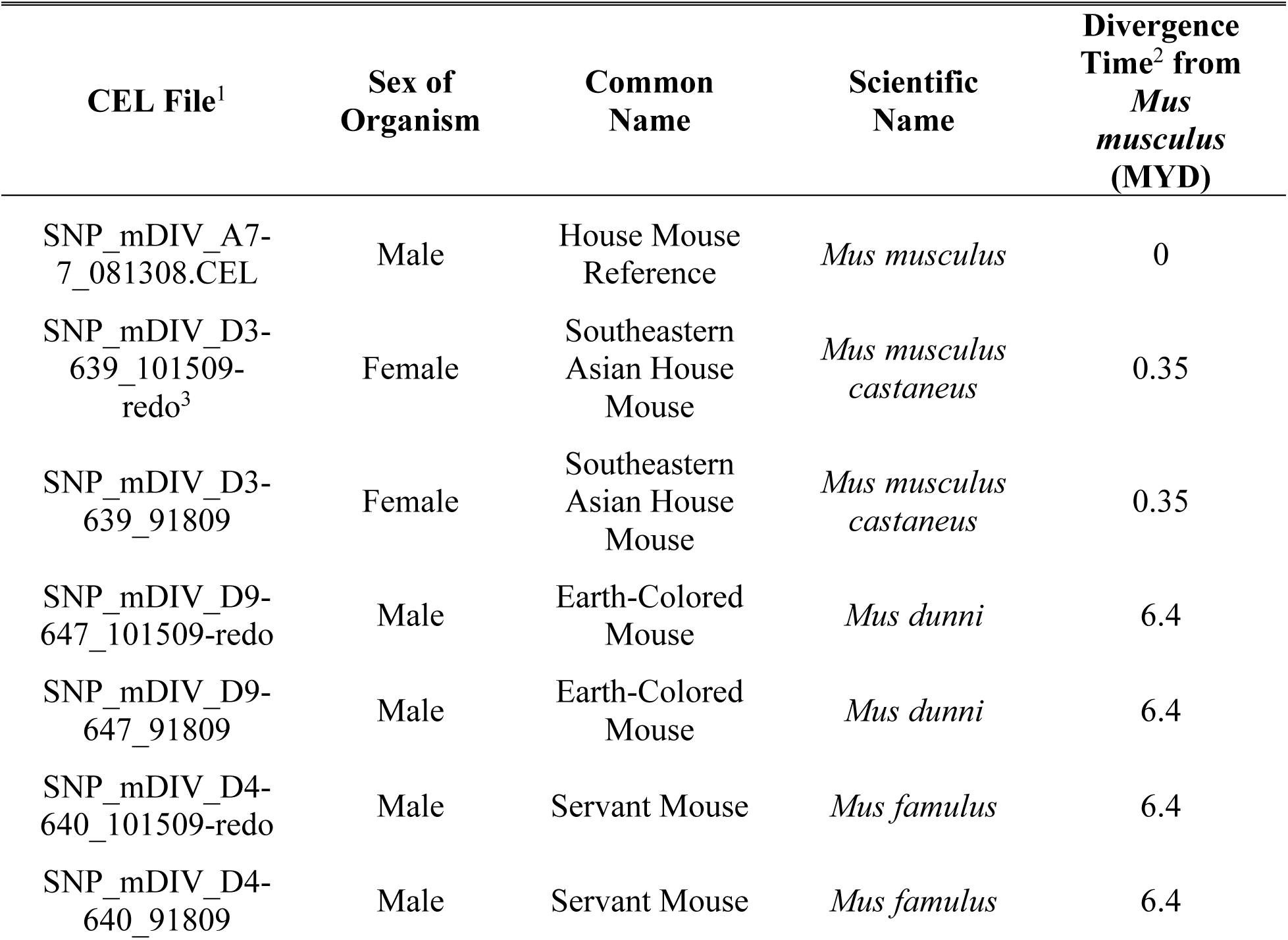
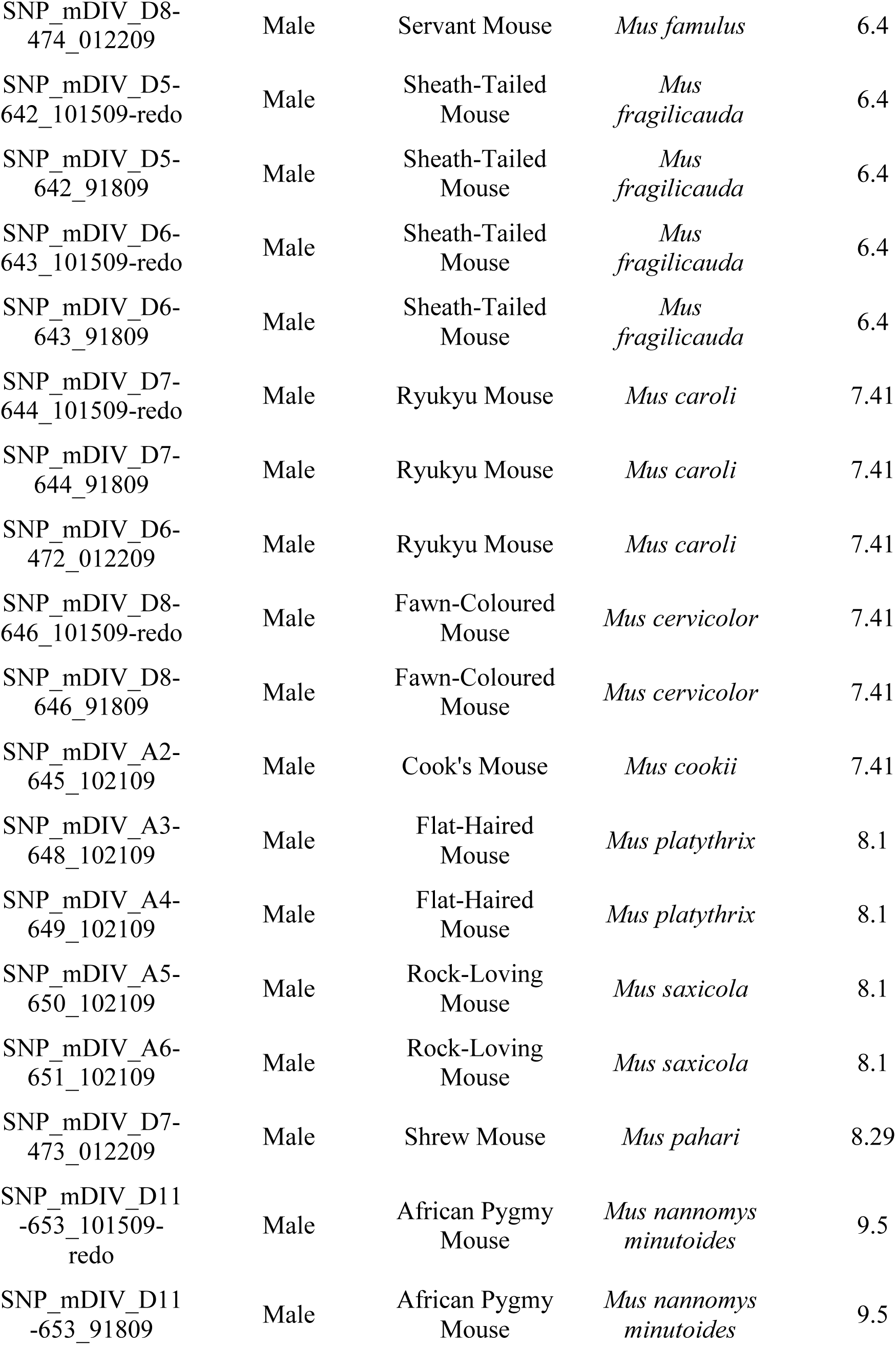
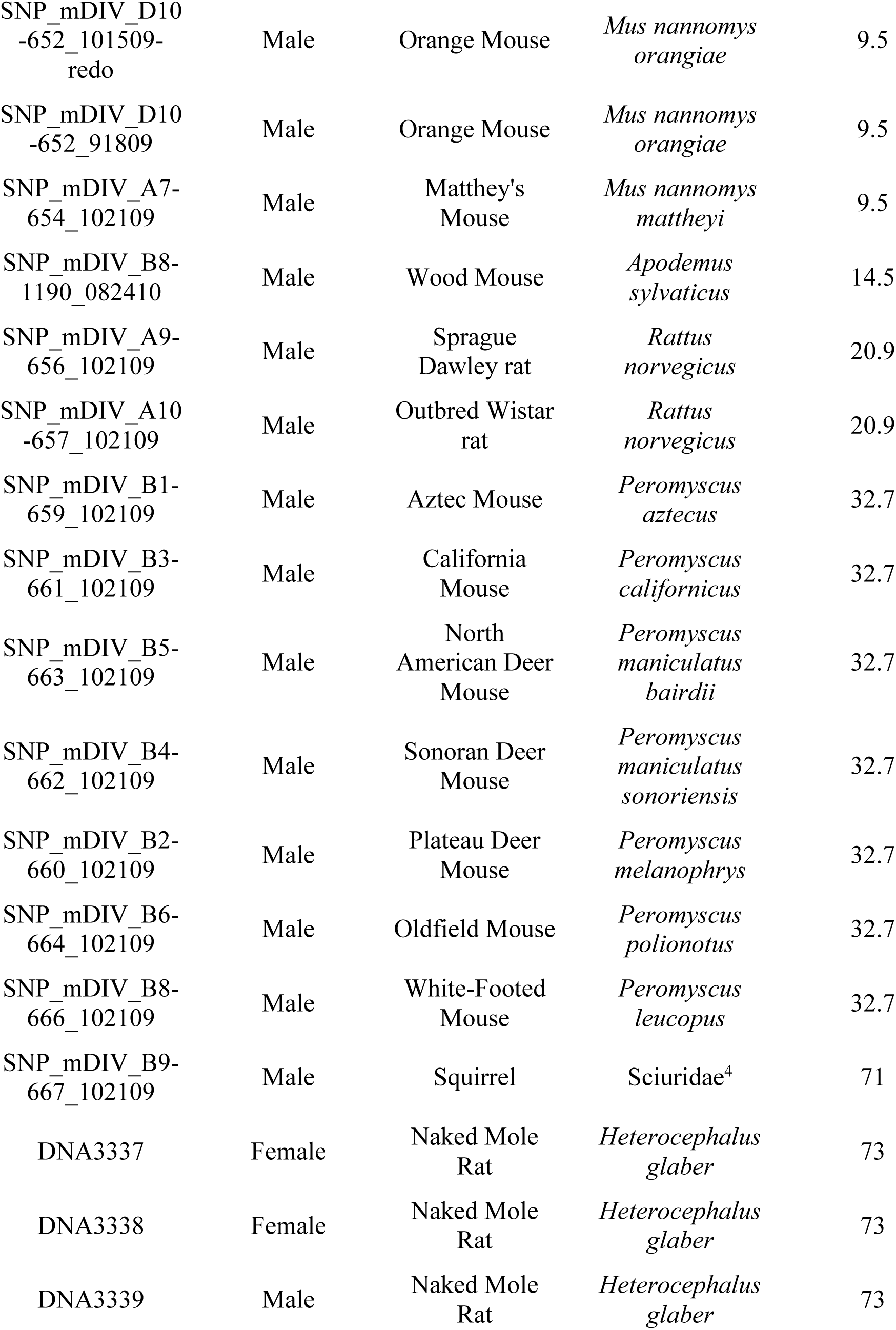
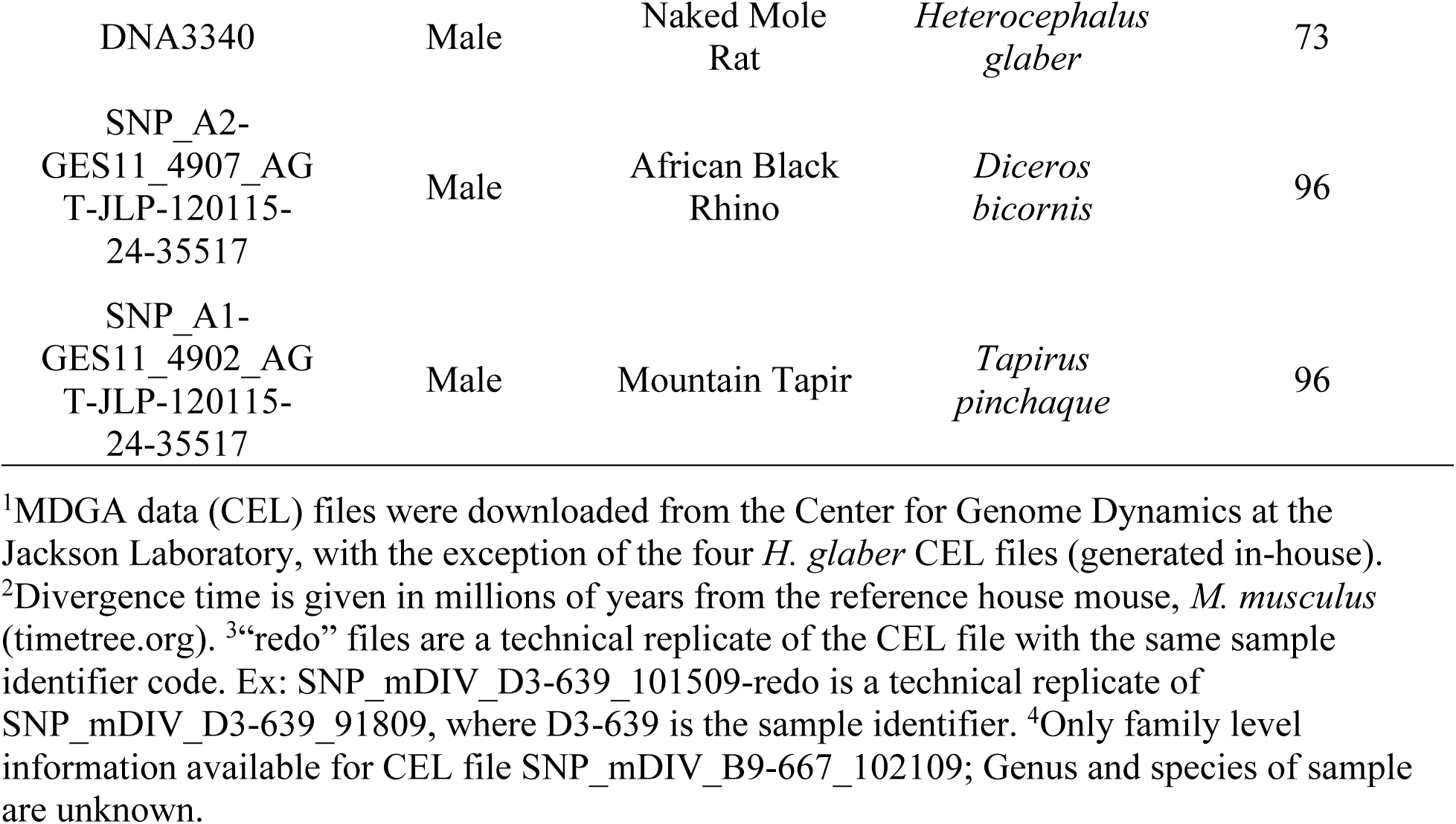
Forty-four MDGA data (CEL) files of the present study.

| CEL File<sup>1</sup> | Sex of Organism | Common Name | Scientific Name | Divergence Time<sup>2</sup> from *Mus musculus* (MYD) |
| --- | --- | --- | --- | --- |
| SNP_mDIV_A7-7_081308.CEL | Male | House Mouse Reference | *Mus musculus* | 0 |
| SNP_mDIV_D3-639_101509-redo<sup>3</sup> | Female | Southeastern Asian House Mouse | *Mus musculus castaneus* | 0.35 |
| SNP_mDIV_D3-639_91809 | Female | Southeastern Asian House Mouse | *Mus musculus castaneus* | 0.35 |
| SNP_mDIV_D9-647_101509-redo | Male | Earth-Colored Mouse | *Mus dunni* | 6.4 |
| SNP_mDIV_D9-647_91809 | Male | Earth-Colored Mouse | *Mus dunni* | 6.4 |
| SNP_mDIV_D4-640_101509-redo | Male | Servant Mouse | *Mus famulus* | 6.4 |
| SNP_mDIV_D4-640_91809 | Male | Servant Mouse | *Mus famulus* | 6.4 |
| SNP_mDIV_D8-474_012209 | Male | Servant Mouse | *Mus famulus* | 6.4 |
| SNP_mDIV_D5-642_101509-redo | Male | Sheath-Tailed Mouse | *Mus fragilicauda* | 6.4 |
| SNP_mDIV_D5-642_91809 | Male | Sheath-Tailed Mouse | *Mus fragilicauda* | 6.4 |
| SNP_mDIV_D6-643_101509-redo | Male | Sheath-Tailed Mouse | *Mus fragilicauda* | 6.4 |
| SNP_mDIV_D6-643_91809 | Male | Sheath-Tailed Mouse | *Mus fragilicauda* | 6.4 |
| SNP_mDIV_D7-644_101509-redo | Male | Ryukyu Mouse | *Mus caroli* | 7.41 |
| SNP_mDIV_D7-644_91809 | Male | Ryukyu Mouse | *Mus caroli* | 7.41 |
| SNP_mDIV_D6-472_012209 | Male | Ryukyu Mouse | *Mus caroli* | 7.41 |
| SNP_mDIV_D8-646_101509-redo | Male | Fawn-Coloured Mouse | *Mus cervicolor* | 7.41 |
| SNP_mDIV_D8-646_91809 | Male | Fawn-Coloured Mouse | *Mus cervicolor* | 7.41 |
| SNP_mDIV_A2-645_102109 | Male | Cook's Mouse | *Mus cookii* | 7.41 |
| SNP_mDIV_A3-648_102109 | Male | Flat-Haired Mouse | *Mus platythrix* | 8.1 |
| SNP_mDIV_A4-649_102109 | Male | Flat-Haired Mouse | *Mus platythrix* | 8.1 |
| SNP_mDIV_A5-650_102109 | Male | Rock-Loving Mouse | *Mus saxicola* | 8.1 |
| SNP_mDIV_A6-651_102109 | Male | Rock-Loving Mouse | *Mus saxicola* | 8.1 |
| SNP_mDIV_D7-473_012209 | Male | Shrew Mouse | *Mus pahari* | 8.29 |
| SNP_mDIV_D11-653_101509-redo | Male | African Pygmy Mouse | *Mus nannomys minutoides* | 9.5 |
| SNP_mDIV_D11-653_91809 | Male | African Pygmy Mouse | *Mus nannomys minutoides* | 9.5 |
| *Mouse single nucleotide polymorphic targets for cross hybridization in rodents* |  |  |  |  |
| SNP_mDIV_D10<br>-652_101509-redo | Male | Orange Mouse | *Mus nannomys*<br>*orangiae* | 9.5 |
| SNP_mDIV_D10<br>-652_91809 | Male | Orange Mouse | *Mus nannomys*<br>*orangiae* | 9.5 |
| SNP_mDIV_A7-654_102109 | Male | Matthey's<br>Mouse | *Mus nannomys*<br>*mattheyi* | 9.5 |
| SNP_mDIV_B8-1190_082410 | Male | Wood Mouse | *Apodemus*<br>*sylvaticus* | 14.5 |
| SNP_mDIV_A9-656_102109 | Male | Sprague<br>Dawley rat | *Rattus*<br>*norvegicus* | 20.9 |
| SNP_mDIV_A10<br>-657_102109 | Male | Outbred Wistar<br>rat | *Rattus*<br>*norvegicus* | 20.9 |
| SNP_mDIV_B1-659_102109 | Male | Aztec Mouse | *Peromyscus*<br>*aztecus* | 32.7 |
| SNP_mDIV_B3-661_102109 | Male | California<br>Mouse | *Peromyscus*<br>*californicus* | 32.7 |
| SNP_mDIV_B5-663_102109 | Male | North<br>American Deer<br>Mouse | *Peromyscus*<br>*maniculatus*<br>*bairdii* | 32.7 |
| SNP_mDIV_B4-662_102109 | Male | Sonoran Deer<br>Mouse | *Peromyscus*<br>*maniculatus*<br>*sonoriensis* | 32.7 |
| SNP_mDIV_B2-660_102109 | Male | Plateau Deer<br>Mouse | *Peromyscus*<br>*melanophrys* | 32.7 |
| SNP_mDIV_B6-664_102109 | Male | Oldfield Mouse | *Peromyscus*<br>*polionotus* | 32.7 |
| SNP_mDIV_B8-666_102109 | Male | White-Footed<br>Mouse | *Peromyscus*<br>*leucopus* | 32.7 |
| SNP_mDIV_B9-667_102109 | Male | Squirrel | Sciuridae<sup>4</sup> | 71 |
| DNA3337 | Female | Naked Mole<br>Rat | *Heterocephalus*<br>*glaber* | 73 |
| DNA3338 | Female | Naked Mole<br>Rat | *Heterocephalus*<br>*glaber* | 73 |
| DNA3339 | Male | Naked Mole<br>Rat | *Heterocephalus*<br>*glaber* | 73 |
| **Mouse single nucleotide polymorphic targets for cross hybridization in rodents** |  |  |  |  |
| DNA3340 | Male | Naked Mole Rat | *Heterocephalus glaber* | 73 |
| SNP_A2-GES11_4907_AG<br>T-JLP-120115-24-35517 | Male | African Black Rhino | *Diceros bicornis* | 96 |
| SNP_A1-GES11_4902_AG<br>T-JLP-120115-24-35517 | Male | Mountain Tapir | *Tapirus pinchaque* | 96 |

<sup>1</sup>MDGA data (CEL) files were downloaded from the Center for Genome Dynamics at the Jackson Laboratory, with the exception of the four *H. glaber* CEL files (generated in-house).
<sup>2</sup>Divergence time is given in millions of years from the reference house mouse, *M. musculus* (timetree.org). <sup>3</sup>“redo” files are a technical replicate of the CEL file with the same sample identifier code. Ex: SNP\_mDIV\_D3-639\_101509-redo is a technical replicate of SNP\_mDIV\_D3-639\_91809, where D3-639 is the sample identifier. <sup>4</sup>Only family level information available for CEL file SNP\_mDIV\_B9-667\_102109; Genus and species of sample are unknown.

**S2 Table..**
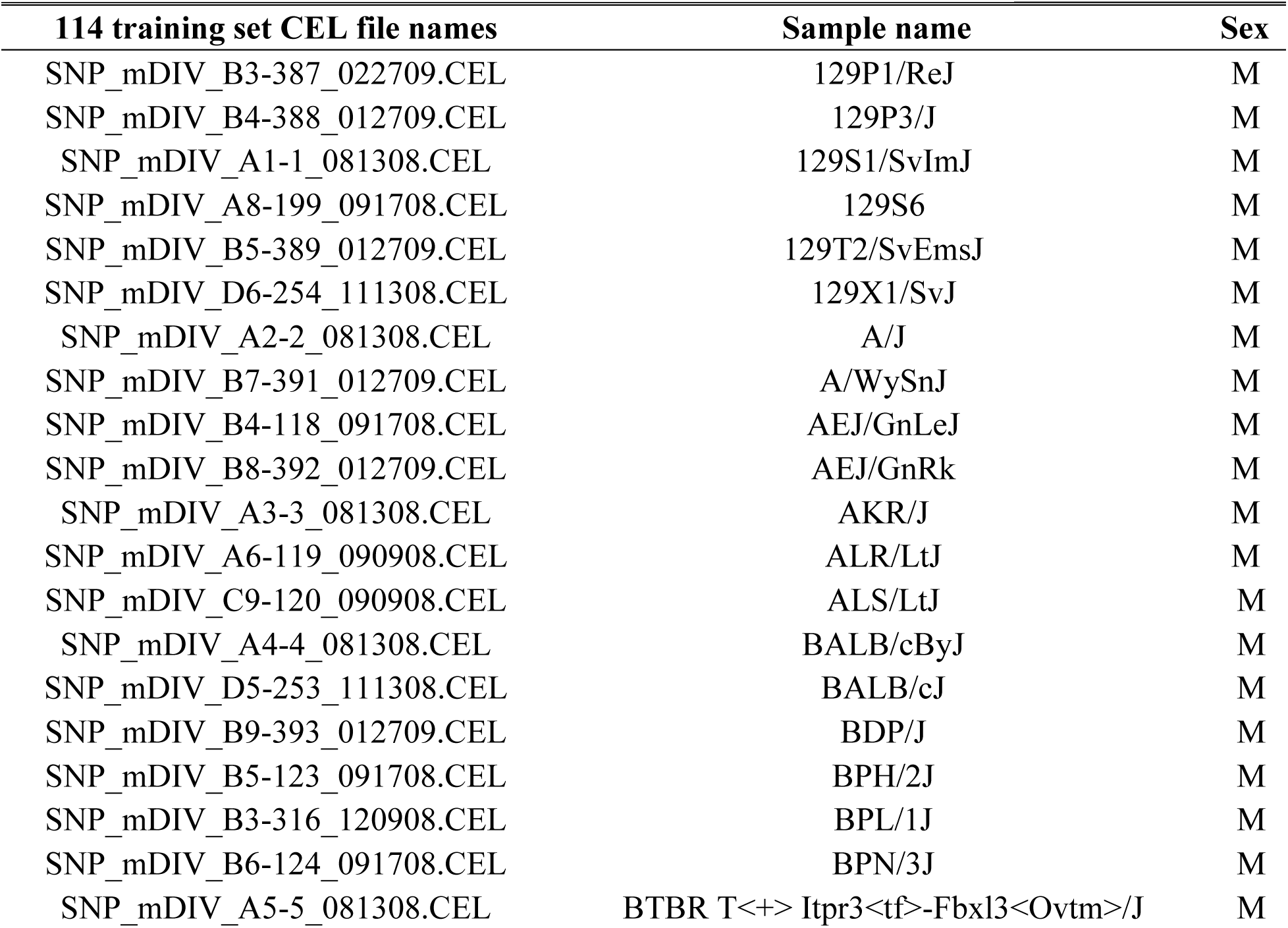
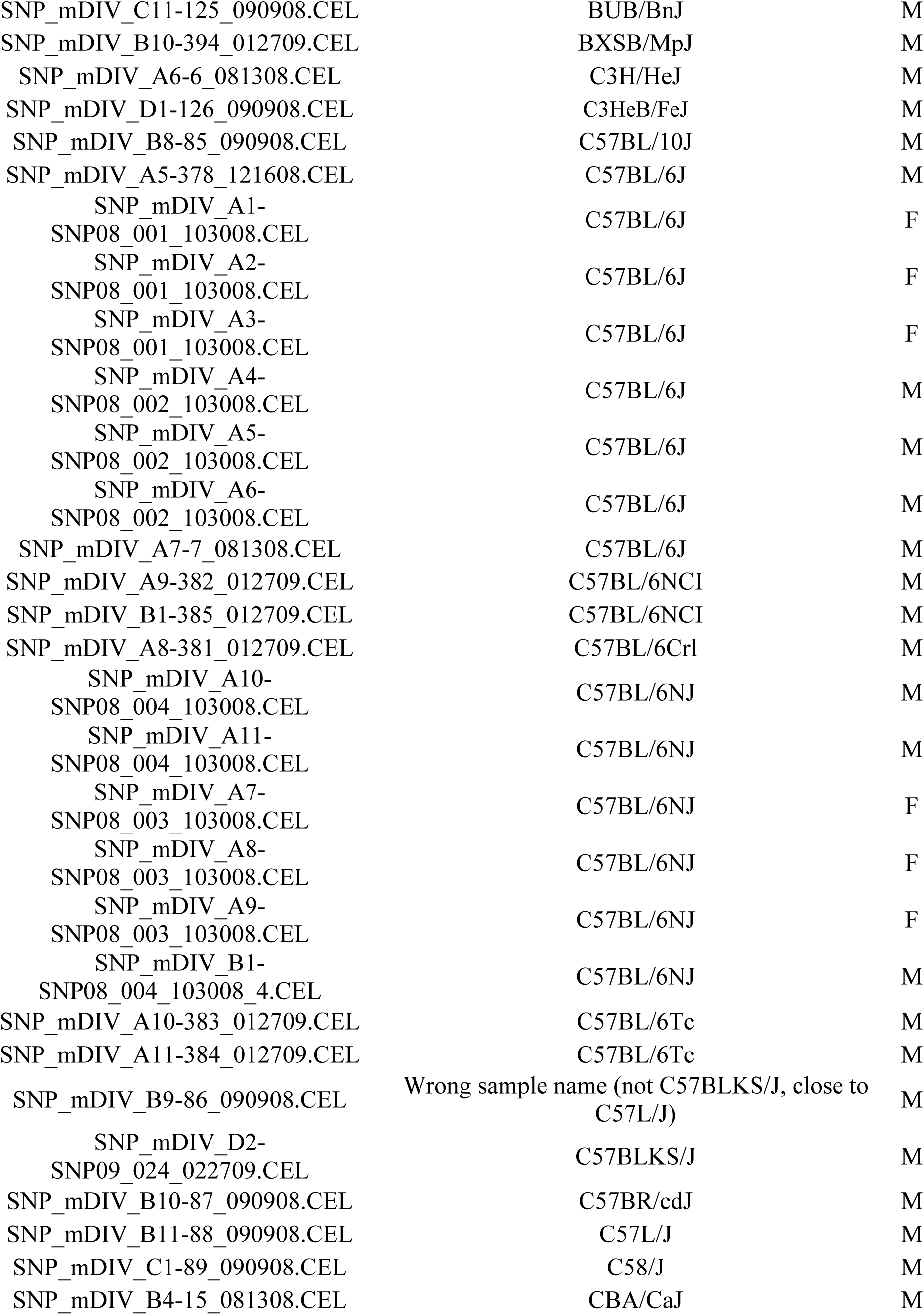
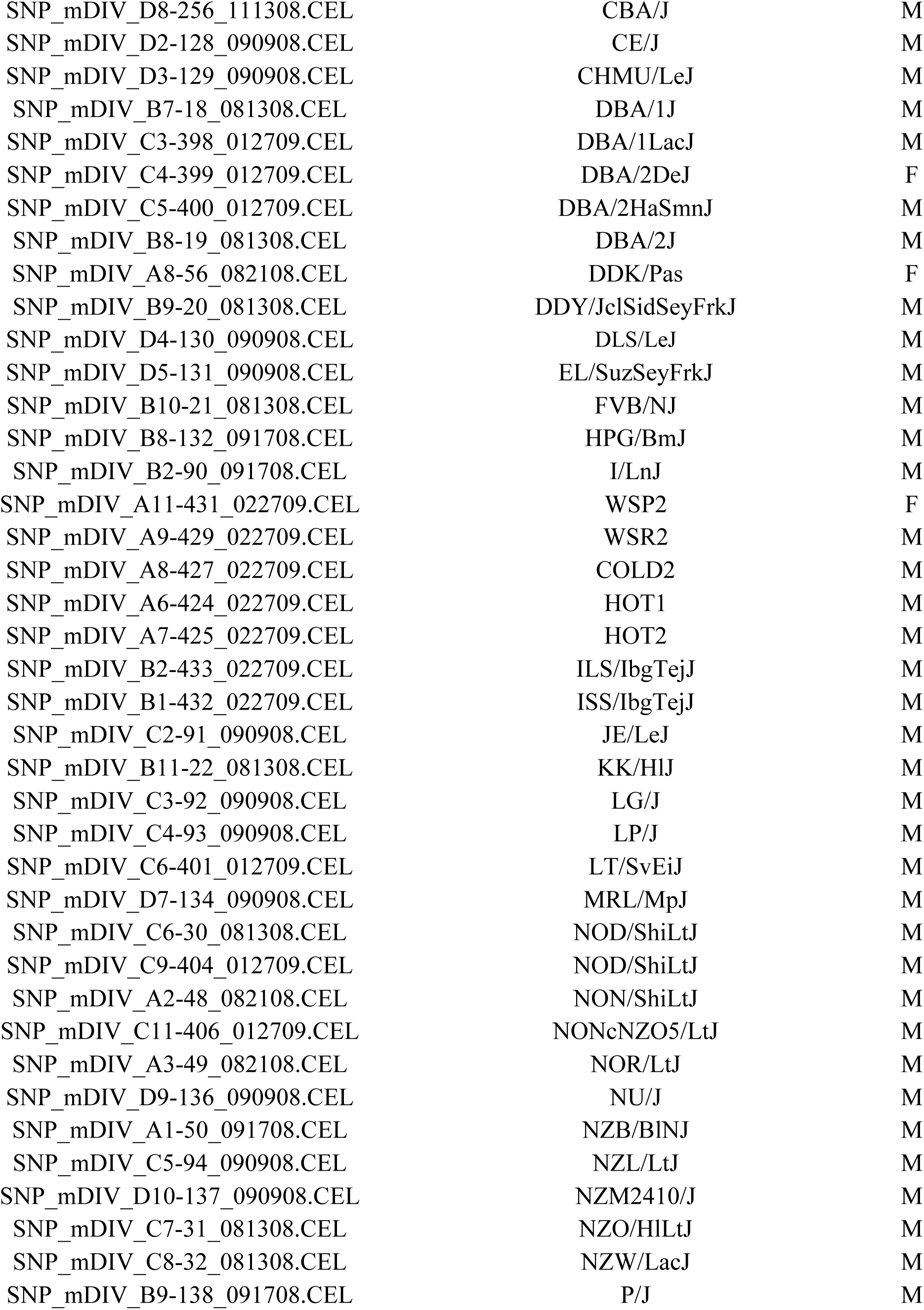
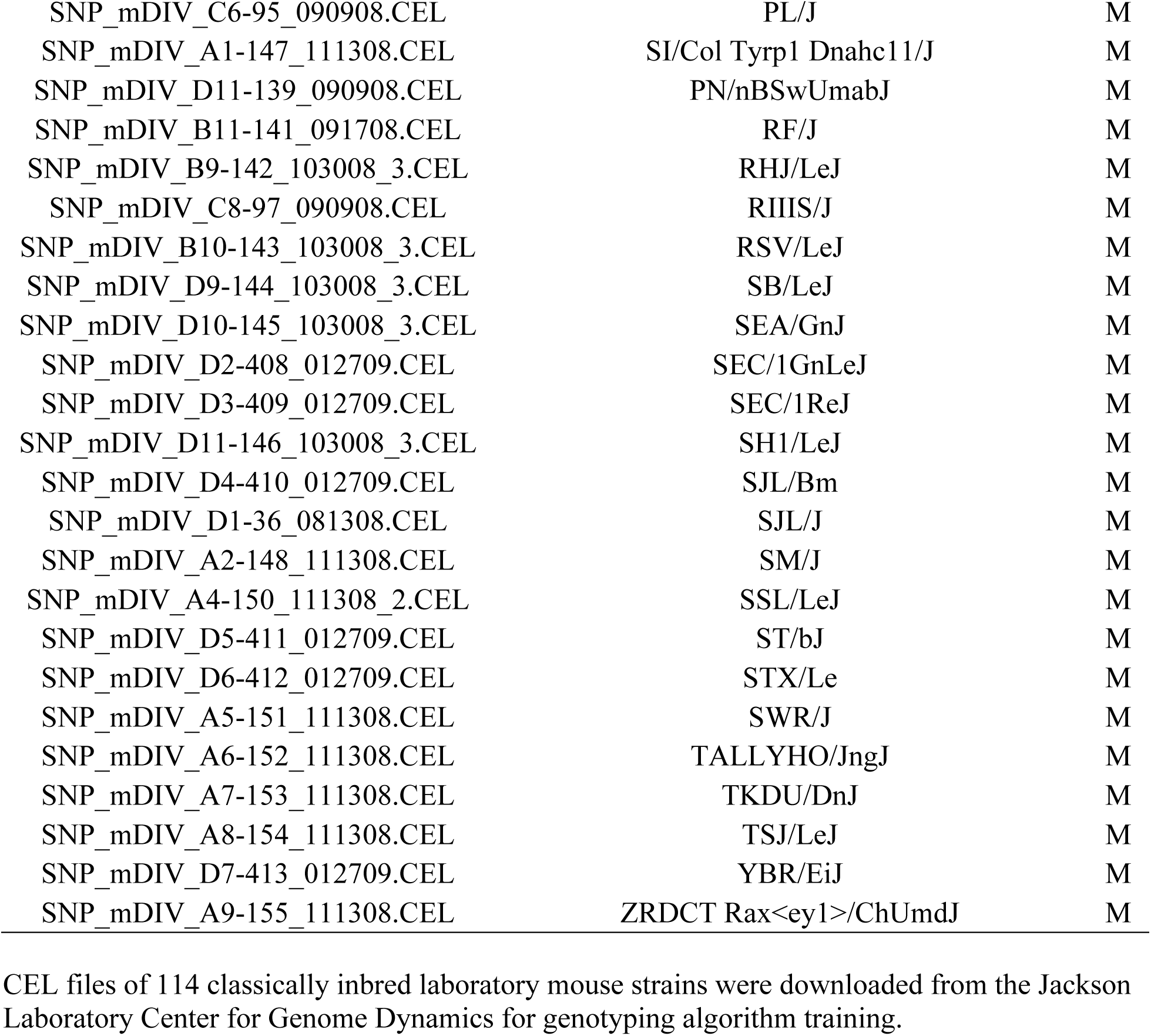
Training set of samples for genotyping algorithm (n=114).

**S3 Table. Genotype summary of 44 study samples genotyped at 493,290 single nucleotide polymorphic loci located across the genome of *Mus musculus*.** Genotyping summary results for all 44 Mouse Diversity Genotyping Array data files. **(Too large to display. Please see separate PDF file)**

**S4 Table..** Study species evaluated with publicly available nuclear genome sequence information.

| Sample Name | Scientific Name | Newest Assembly |
| --- | --- | --- |
| House Mouse | *Mus musculus* | GRCm38.p6 |
| Ryukyu Mouse | *Mus caroli* | CAROLI_EIJ_v1.1 |
| Gairdner's Shrewmouse | *Mus pahari* | PAHARI_EIJ_v1.1 |
| Sprague Dawley Rat | *Rattus norvegicus* | Rnor_6.0 |
| North American Deer Mouse | *Peromyscus maniculatus* | Pman_1.0 |

Genomes accessed through the NCBI Genomes FTP site of samples under study (<ftp.ncbi.nlm.nih.gov/genomes/>).

